# A Single, Reproducible Dimension Organizes Associations Between the Early Environment and the Developing Brain

**DOI:** 10.64898/2026.09.26.754719

**Authors:** Chase Antonacci, Kaitlyn Kwan, Eugenia Giampetruzzi, Jessica P. Uy, Ian H. Gotlib

## Abstract

Childhood adversity has enduring consequences for health and development across the lifespan, yet the organizing structure linking diverse environmental experiences to the developing brain remains poorly understood. We jointly modeled 104 measures of children’s environments, spanning interpersonal, household, and neighborhood contexts, with 7 neuroimaging modalities of brain structure and function in 5,976 youth. Despite this breadth, only a single adversity-brain dimension emerged and replicated in held-out data. This dimension was defined primarily by socioeconomic and neighborhood deprivation and selectively expressed in lower cortical surface area and distributed functional connectivity concentrated in sensorimotor systems. Total cortical surface area reproduced 96% of the regional association, indicating a global rather than regionally specific structural signature. This dimension predicted poorer cognition and cardiometabolic health six years later. These findings reveal a strikingly selective, low-dimensional relation between children’s environments and neurodevelopment, with socioeconomic deprivation emerging as the principal environmental axis associated with the developing brain.

## Introduction

Early adversity is among the strongest environmental predictors of cognition and mental and physical health across the lifespan^1–3^. Adverse experiences, from socioeconomic disadvantage and neighborhood disorder to family instability, caregiver psychopathology, and violence, frequently co-occur and are associated with greater risk for psychopathology and poorer cognitive and educational outcomes^1,4,5^. Because childhood and adolescence are periods of substantial neurodevelopmental change and experience-dependent plasticity^6,7^, the developing brain has been proposed as a central pathway through which individual differences in the early environment become linked to later functioning^8,9^. Consistent with this possibility, childhood stress and disadvantage have been related to variation in cortical morphology, white-matter organization, task-evoked activation, and the functional architecture of large-scale brain networks^10–12^. The extent to which these reported associations reflect a coherent underlying structure, however, is not clear. The many associations reported across exposures and imaging phenotypes could reflect numerous distinct effects of particular experiences on particular neural systems or, alternatively, a much smaller number of organizing dimensions linking variation in the early environment to neurodevelopment. Distinguishing between these possibilities is critical for identifying which features of children’s environments are most consequential for neurodevelopment and whether the extensive neural correlates of adversity reported across the literature represent distinct biological processes or manifestations of a common underlying structure.

Addressing this question is complicated by substantial intercorrelation among adverse experiences^13,14^, which can obscure meaningful distinctions among environmental exposures. Dimensional frameworks have proposed that qualitatively distinct experiences, most prominently threat and deprivation and, in other models, environmental unpredictability, influence behavioral and psychobiological development through partially distinct pathways^15–17^. Other approaches emphasize cumulative adversity^18,19^ or identify alternative latent structures from patterns of co-occurring exposures^20,21^. Although these models make different claims about which features of experience should organize developmental variation, evidence concerning the role of brain metrics has generally come from studies in which theoretically defined exposures or dimensions are examined separately. While such designs can establish associations with particular neural phenotypes, they cannot determine whether proposed distinctions among adverse experiences continue to be evident when correlated exposures are considered together. Consequently, it remains unclear whether the distinctions among adverse experiences proposed by current frameworks actually organize variation in neurodevelopment.

Importantly, the neural correlates of adversity have been found to be distributed across a wide range of structural and functional phenotypes. Adversity has been associated with cortical morphology, white matter microstructure, intrinsic functional connectivity, and task-evoked activation, although these phenotypes are rarely evaluated within the same analysis^8,10–12^. Consequently, an adversity-brain association observed within one imaging modality provides little information about its prominence or importance relative to associations elsewhere in the brain. A smaller literature has begun to address this problem using multivariate methods relating high-dimensional phenotypic data to brain organization. For example, using canonical correlation analysis and related approaches, researchers identified a single dominant “positive-negative” mode linking functional connectivity to a set of demographic and behavioral traits^22^. More recent multimodal analyses in adolescents recovered richer sets of ‘modes,’ including patterns dominated by perinatal risk and socioeconomic stratification^23^ and 14 reliable modes spanning 4 imaging modalities^24^. Although these studies demonstrated that shared dimensions can be recovered from complex behavioral and neuroimaging data, they were not designed to determine how a comprehensive representation of childhood adversity maps onto the brain. Environmental exposures have generally been embedded within broader batteries containing cognition, behavior, or psychopathology, leaving unresolved the question of which features of adverse experience account for the resulting brain associations.

Therefore, a central challenge in adversity neuroscience is to identify which environment-brain associations remain when variation across both domains is modeled simultaneously. Associations identified from individual exposures or single imaging modalities may reflect variance shared among correlated measures, making it difficult to determine whether reported findings represent distinct and reproducible features of adversity-related neurodevelopment or reflect a smaller number of shared environment-brain associations. Addressing this problem requires modeling correlated environmental exposures together while simultaneously allowing multiple structural and functional neural phenotypes to contribute without specifying *a priori* where in the brain adversity-related differences should occur. Importantly, any resulting organization must also generalize beyond the sample in which it was estimated, particularly because brain-wide associations are typically small and can be substantially inflated at conventional sample sizes^25,26^. Joint modeling with out-of-sample validation can establish both which distinctions among adverse experiences are reproducibly associated with neurodevelopment and where in the brain those associations are expressed most consistently.

The present study was designed to examine whether the many reported associations between early adversity and the brain reduce to a smaller number of reproducible environment-brain dimensions and, if so, which environmental exposures and neural phenotypes define them. To address this question, we used sparse generalized canonical correlation analysis (SGCCA), which identifies weighted combinations of features that maximize shared covariance across multiple high-dimensional datasets. Unlike approaches that characterize the structure of environmental exposures independent of their neural correlates, SGCCA allows the organization of the environmental and neural data to be estimated jointly, without imposing a theoretical taxonomy of adversity or designating a specific neural system *a priori*. Applying this approach in the Adolescent Brain Cognitive Development (ABCD) Study, we related 104 environmental measures spanning interpersonal, household, and neighborhood contexts to 7 measures of brain structure and function: cortical surface area (SA), cortical thickness (CT), white-matter fractional anisotropy (FA), resting-state functional connectivity (rsFC), and activation during the Monetary Incentive Delay (MID), Stop-Signal (SST), and Emotional N-Back (ENB) tasks. We estimated the multivariate structure in 70% of families and applied the resulting weights unchanged to the remaining 30%, retaining only dimensions that replicated out of sample. We then tested whether the resulting dimensions predicted cognition, psychopathology, and cardiometabolic health six years later and whether the identified brain features contributed predictive information beyond that provide by adversity alone. Together, these analyses allowed us to examine the dimensionality, neurobiological specificity, and prospective significance of reproducible relations between the early environment and the developing brain.

## Results

### Participant Characteristics

Exposure to adversity was widely but modestly endorsed in the ABCD cohort (Table 1). 86.6% of caregivers reported at least one of 25 stressful life events (*M*=3.11, *SD*=2.51) and 82.5% reported at least one of nine family-conflict items (*M*=2.44, *SD*=1.94); 34.4% of youth had experienced at least one of the 18 KSADS-screener traumatic events (9.0% two or more). At the year-6 follow-up, participants reported mild to moderate symptoms of psychopathology (*M*_CBCL Total_=12.9±15.4) and NIH-TB cognition scores (*M*_Crystallized_= 106.7±15.3, *M*_Fluid_=114.3±18.9) and cardiometabolic markers within typical ranges (*M*_waist-to-height_=0.48±0.07; *M*_systolic BP_=111.5±11.6 mmHg; *M*_diastolic BP_=63.8±8.6 mmHg; *M*_total cholesterol_=152.7±28.7 mg/dL; *M*_glycated hemoglobin_=5.38%±0.30). Relative to the 5,884 baseline participants without usable imaging data, the analytic sample (n=5,976) was more likely to be White (59.0% vs. 45.2%, *p*<.0001) and to report household income >$100,000 (48.2% vs. 35.5%, *p*<.0001), a selection pattern consistent with documented associations between scan quality and sociodemographic factors^27^. The training and held-out samples did not differ significantly on sociodemographic characteristics or the primary study variables (Table 1).

**Table 1:**
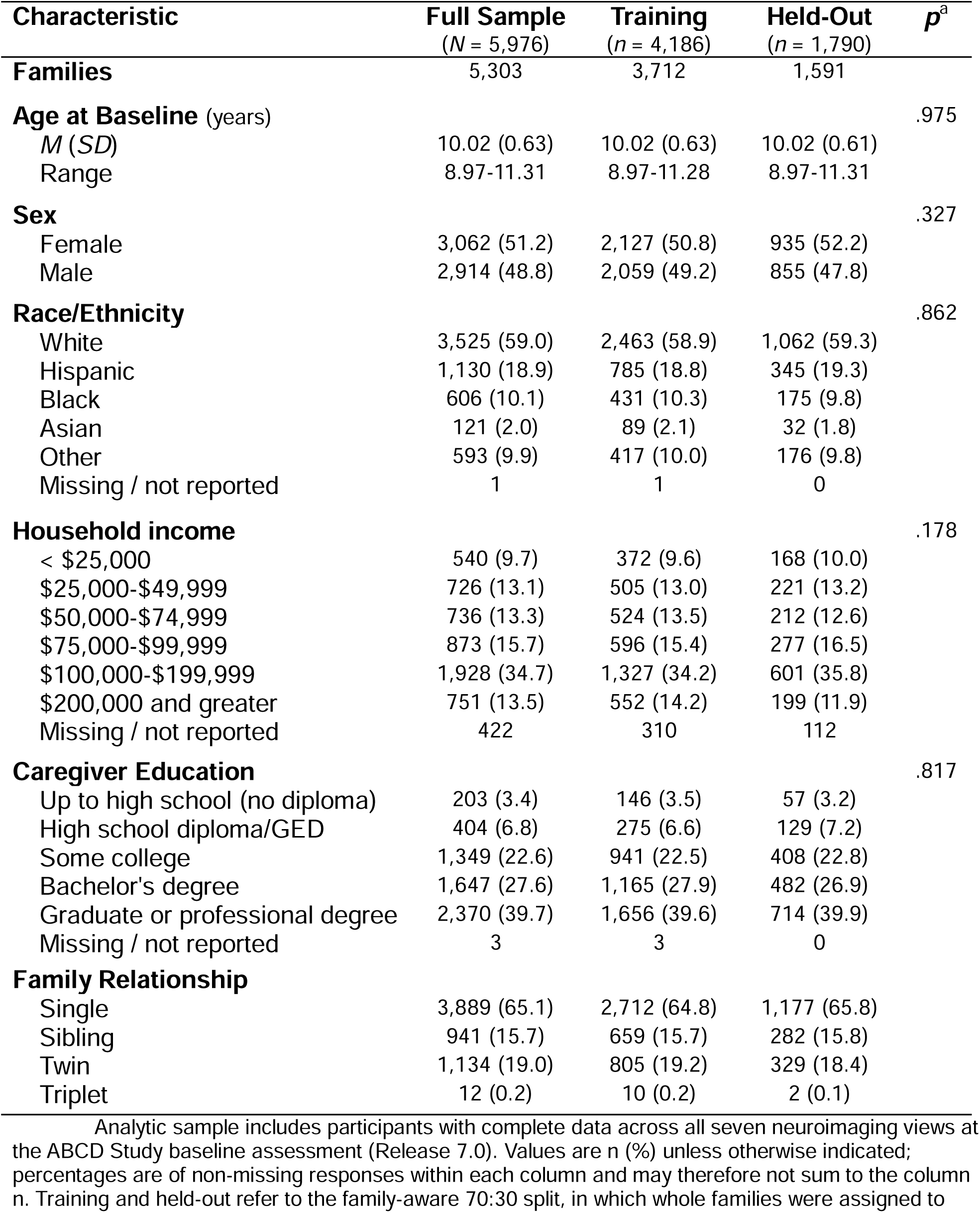
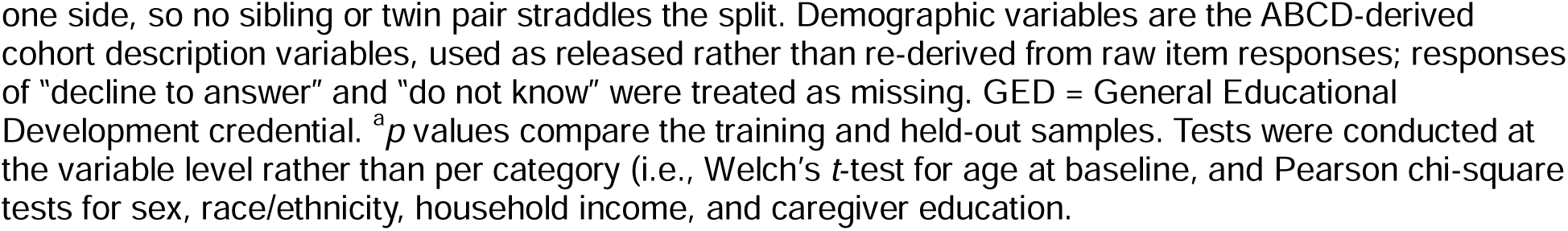
Sample Characteristics.

Analytic sample includes participants with complete data across all seven neuroimaging views at the ABCD Study baseline assessment (Release 7.0).
| Characteristic | Full Sample<br>(N = 5,976) | Training<br>(n = 4,186) | Held-Out<br>(n = 1,790) | *p*<sup>a</sup> |
| --- | --- | --- | --- | --- |
| **Families** | 5,303 | 3,712 | 1,591 |  |
| **Age at Baseline** (years) |  |  |  | .975 |
| *M* (*SD*) | 10.02 (0.63) | 10.02 (0.63) | 10.02 (0.61) |  |
| Range | 8.97-11.31 | 8.97-11.28 | 8.97-11.31 |  |
| **Sex** |  |  |  | .327 |
| Female | 3,062 (51.2) | 2,127 (50.8) | 935 (52.2) |  |
| Male | 2,914 (48.8) | 2,059 (49.2) | 855 (47.8) |  |
| **Race/Ethnicity** |  |  |  | .862 |
| White | 3,525 (59.0) | 2,463 (58.9) | 1,062 (59.3) |  |
| Hispanic | 1,130 (18.9) | 785 (18.8) | 345 (19.3) |  |
| Black | 606 (10.1) | 431 (10.3) | 175 (9.8) |  |
| Asian | 121 (2.0) | 89 (2.1) | 32 (1.8) |  |
| Other | 593 (9.9) | 417 (10.0) | 176 (9.8) |  |
| Missing / not reported | 1 | 1 | 0 |  |
| **Household income** |  |  |  | .178 |
| < \$25,000 | 540 (9.7) | 372 (9.6) | 168 (10.0) | |
| \$25,000-\$49,999 | 726 (13.1) | 505 (13.0) | 221 (13.2) | |
| \$50,000-\$74,999 | 736 (13.3) | 524 (13.5) | 212 (12.6) | |
| \$75,000-\$99,999 | 873 (15.7) | 596 (15.4) | 277 (16.5) | |
| \$100,000-\$199,999 | 1,928 (34.7) | 1,327 (34.2) | 601 (35.8) | |
| \$200,000 and greater | 751 (13.5) | 552 (14.2) | 199 (11.9) | |
| Missing / not reported | 422 | 310 | 112 |  |
| **Caregiver Education** |  |  |  | .817 |
| Up to high school (no diploma) | 203 (3.4) | 146 (3.5) | 57 (3.2) |  |
| High school diploma/GED | 404 (6.8) | 275 (6.6) | 129 (7.2) |  |
| Some college | 1,349 (22.6) | 941 (22.5) | 408 (22.8) |  |
| Bachelor's degree | 1,647 (27.6) | 1,165 (27.9) | 482 (26.9) |  |
| Graduate or professional degree | 2,370 (39.7) | 1,656 (39.6) | 714 (39.9) |  |
| Missing / not reported | 3 | 3 | 0 |  |
| **Family Relationship** |  |  |  |  |
| Single | 3,889 (65.1) | 2,712 (64.8) | 1,177 (65.8) |  |
| Sibling | 941 (15.7) | 659 (15.7) | 282 (15.8) |  |
| Twin | 1,134 (19.0) | 805 (19.2) | 329 (18.4) |  |
| Triplet | 12 (0.2) | 10 (0.2) | 2 (0.1) |  |

one side, so no sibling or twin pair straddles the split. Demographic variables are the ABCD-derived cohort description variables, used as released rather than re-derived from raw item responses; responses of “decline to answer” and “do not know” were treated as missing. GED = General Educational Development credential. <sup>a</sup>*p* values compare the training and held-out samples. Tests were conducted at the variable level rather than per category (i.e., Welch’s *t*-test for age at baseline, and Pearson chi-square tests for sex, race/ethnicity, household income, and caregiver education).

### A Single, Cross-Validated Adversity-Brain Dimension

We first tested whether the early environment, measured across interpersonal, household, and neighborhood levels, maps onto a reproducible pattern of brain structure and function. We fit a sparse generalized CCA model relating the 104-feature adversity view to all 7 imaging views (503 features) in the training set and extracted 5 components (Fig. 1a). Only the first component replicated in both the training and held-out test sets, exceeding a family-level permutation null in both (mean variance explained across the seven adversity-brain pairs=0.0128 versus null=0.0022; test=0.0125 vs. null=0.0023; both *p*<.001, 10,000 permutations); components 2-5 did not replicate in the held-out set (test *ps*>.47; Fig. 1b). Therefore, we retained a single component and interpret a single adversity-brain axis throughout. Two of the seven imaging views accounted for this axis (Fig. 1c). Cortical SA had the strongest association with the adversity variate (training |*r*|=0.210, *p*<.001; test |*r*|=0.209, *p*=.0003), and resting-state network connectivity was also strong and robust (training |*r*|=0.195, *p*<.001; test |*r*|=0.190, *p*<.001). The remaining five views – CT, white-matter FA, and the three task-fMRI contrasts – had no association with the adversity variate that replicated across both splits (all |*r*|≤0.055). At the subject level, both associations reproduced in held-out families using the non-sparse companion variates (surface area |*r*|=0.226, resting-state |*r*|=0.138; Fig. 1d).

**Figure 1:**
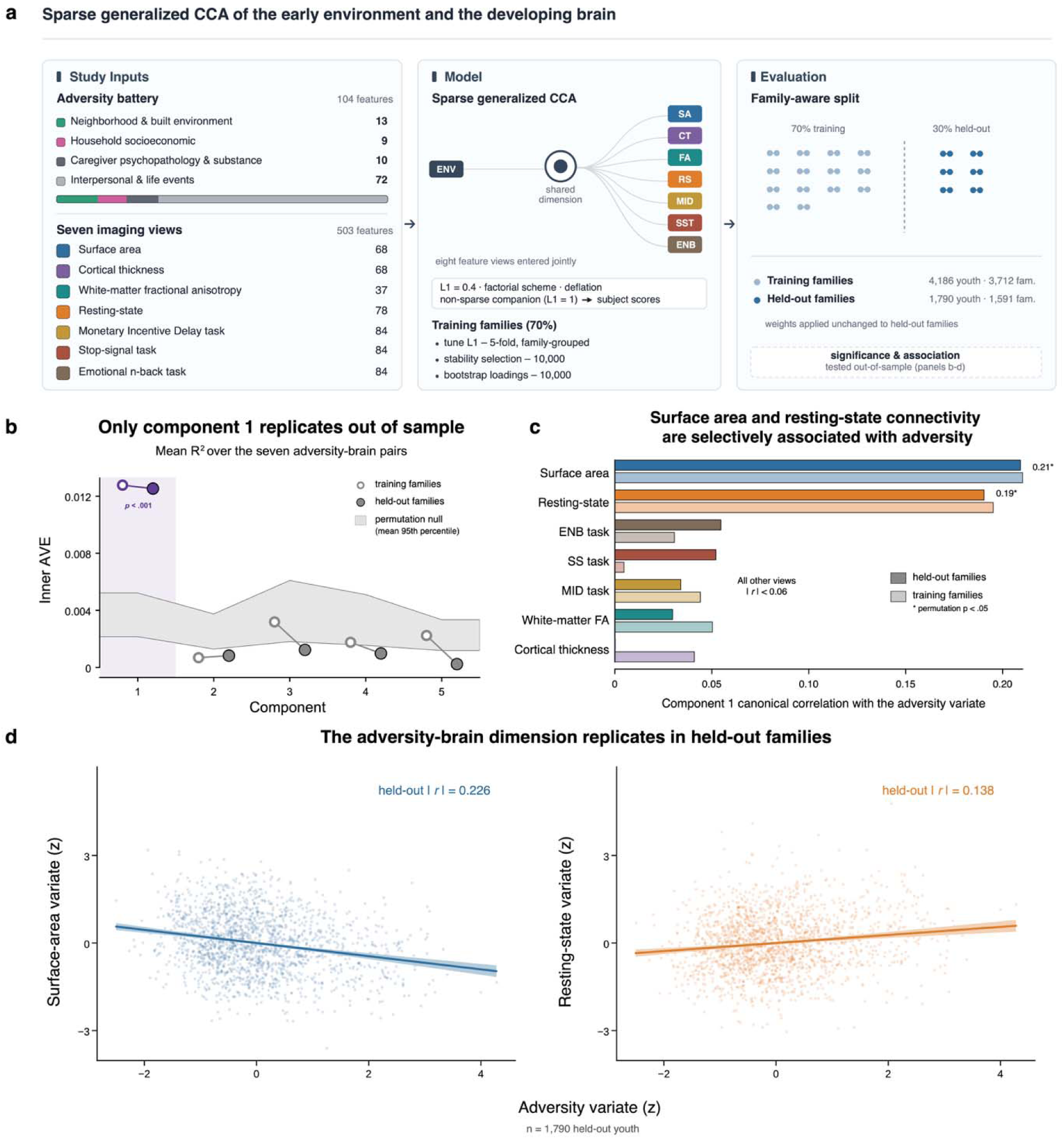
A Single, Reproducible Adversity-Brain Dimension. **(a)** Analytic design. A 104-feature early-adversity battery spanning four domains and seven imaging views were entered jointly into a sparse generalized canonical correlation analysis (L1 = 0.4, factorial scheme, deflation). Weights were estimated in 70% of families (4,186 youth; 3,712 families) using 5-fold family-grouped tuning, stability selection, and bootstrapping (10,000 resamples each) and applied unchanged to held-out families (1,790 youth; 1,591 families). **(b)** Mean inner AVE (average variance explained across the seven adversity-brain view pairs) for each component in training (lighter) and held-out (darker) families, against a permutation null (shaded; mean 95th percentile). Only the first component exceeded the null out of sample (*p* < .001). **(c)** Component-1’s canonical correlation between each imaging view and the adversity variate. Surface area (0.21) and resting-state connectivity (0.19) are the only views reaching significance (permutation *p* < .05); all others |*r*| < 0.06. **(d)** The dimension reproduces in held-out youth, with adversity-variate scores correlating with surface-area (|*r*| = 0.226) and resting-state (|*r*| = 0.138) variate scores (n = 1,790).

With respect to adversity, the first canonical variate was defined principally by socioeconomic and neighborhood deprivation – hereafter the socioeconomic disadvantage dimension. Of the 104 adversity features, 39 received non-zero weights and 19 were stably selected in at least 90% of 10,000 training subsamples (Fig. 2a,b). The largest structural loadings were low child opportunity (*r*=0.734 [0.678, 0.784], *q*=.002), low social mobility (*r*=0.704 [0.648, 0.754], *q*=.001), area deprivation (*r*=0.694 [0.640, 0.742], *q*=.002), and social vulnerability (*r*=0.678 [0.623, 0.726], *q*=.002), all from linked neighborhood data (Fig. 2a). These were followed by more proximal indicators of family socioeconomic resources, including low household income (*r*=0.651 [0.618, 0.681], *q*=.001), low caregiver education (*r*=0.592 [0.553, 0.628], *q*=.002), and caregiver-reported neighborhood danger (*r*=0.506 [0.462, 0.548], *q*=.001). A smaller, second cluster reflected caregiver psychopathology, with the antisocial (*r*=0.362 [0.283, 0.435], *q*=.002) and somatic (*r*=0.353 [0.289, 0.415], *q*=.002) symptom scales stably selected, together with a single interpersonal exposure of witnessing household (interparental) violence (*r*=0.278 [0.219, 0.337], *q*=.001).

**Figure 2:**
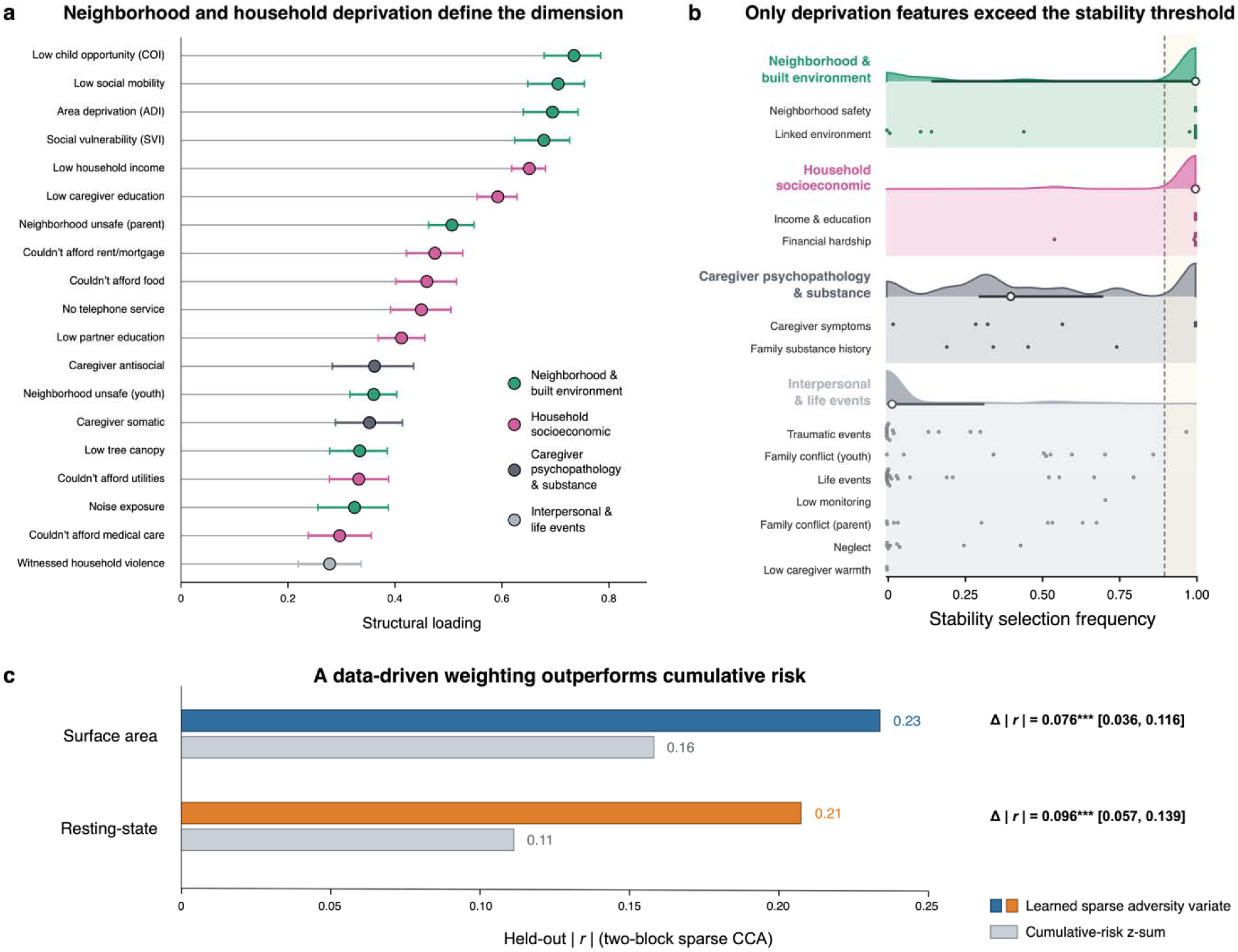
Socioeconomic Disadvantage Defines the Dimension and Outperforms Cumulative-Risk Scoring. **(a)** Bootstrapped loadings (mean ± 95% CI) of individual adversity features, colored by domain. Neighborhood disadvantage (low child opportunity, low social mobility, area deprivation, social vulnerability) and household socioeconomic hardship (low income, low caregiver education) had the largest loadings. **(b)** Stability selection frequency across 10,000 resamples, grouped by domain (clouds show domain-level distributions; points are individual features). Only neighborhood and household deprivation features exceed the 90% threshold (dashed line); caregiver psychopathology and interpersonal/life event features are rarely selected. **(c)** The learned sparse adversity variate predicts held-out brain variates better than an unweighted cumulative risk *z*-sum of the same exposures, for both surface area (|*r*| = 0.23 vs 0.16; Δ|*r*| = 0.076, 95% CI [0.036, 0.116]) and resting-state network functional connectivity (|*r*| 0.21 vs 0.11; Δ|*r*| = 0.096 [0.057, 0.139]). \*\*\**p* < .001.

In total, the 19 stably selected features comprised 6 neighborhood measures, 8 household socioeconomic indicators, both neighborhood-safety scales, two caregiver-psychopathology scales, and one traumatic-event item. No measure of family conflict, neglect, caregiver warmth, parental monitoring, or parent-reported life events met the 90% stability threshold (Fig. 2b). Thus, when interpersonal, household, and neighborhood exposures are modeled jointly, the brain-linked axis most consistently reflects socioeconomic and neighborhood conditions and caregiver psychopathology rather than family conflict, neglect, or parenting behavior. Estimating each modality against adversity independently (two-view models) reproduced these two associations (held-out |*r*|=0.234 surface area, 0.207 resting-state), suggesting they are not artifacts of the joint fit. Notably, this data-driven weighting also predicted the held-out brain variates substantially better than an unweighted cumulative-risk sum of the same 104 exposures (surface area |*r*|=0.23 vs 0.16, Δ|*r*|=0.076 [0.036, 0.116]; resting-state |*r*|=0.21 vs 0.11, Δ|*r*|=0.096 [0.057, 0.139]; Fig. 2c). We decompose these single modality estimates below.

### Global Cortical Surface Area and Distributed Network Connectivity

The above analysis identified a pattern of lower surface area associated with greater adversity but cannot, on its own, determine whether this pattern reflects coordinated differences across specific regions or a global difference in overall cortical size. Therefore, we tested these two possibilities directly. Greater adversity was associated with smaller total cortical SA (i.e., the summed area of all 68 parcels: *β*=-0.202 [-0.224, -0.180], *p*<.0001 in the full sample; *β*=-0.190 [-0.230, -0.150], *p*<.001 in the held-out set). Importantly, substituting this single global measure for the 68 regional features reproduced 96.1% [89.5%, 102.5%] of the out-of-sample regional effect (two-view held-out |*r*|=0.225 vs. 0.234; Fig. 3c) – that is, adversity appears to be associated more strongly with a smaller overall cortex than with focal differences in specific regions. Conversely, residualizing total SA from each parcel reduced the held-out association by more than half (46.7% retained [27.8%, 66.1%] Fig. 3c), and none of the 68 regions remained stably selected (Fig. 3a,b). Thus, once overall size is removed, little regional signal remains. Further, residualizing intracranial volume from each regional feature retained 69.0% [52.9%, 85.2%] of the original association, indicating that the effect is not reducible to head size alone. Consistent with a global effect of adversity on cortical SA, there was little robust regional localization. 12 of 68 regions were stably selected in the unadjusted features, where each region still carries the shared global signal, but none survived once total SA was removed (peak selection=0.78; Fig. 3a,b). Thus, adversity was associated with a smaller cortex overall rather than a spatially specific regional pattern.

**Figure 3:**
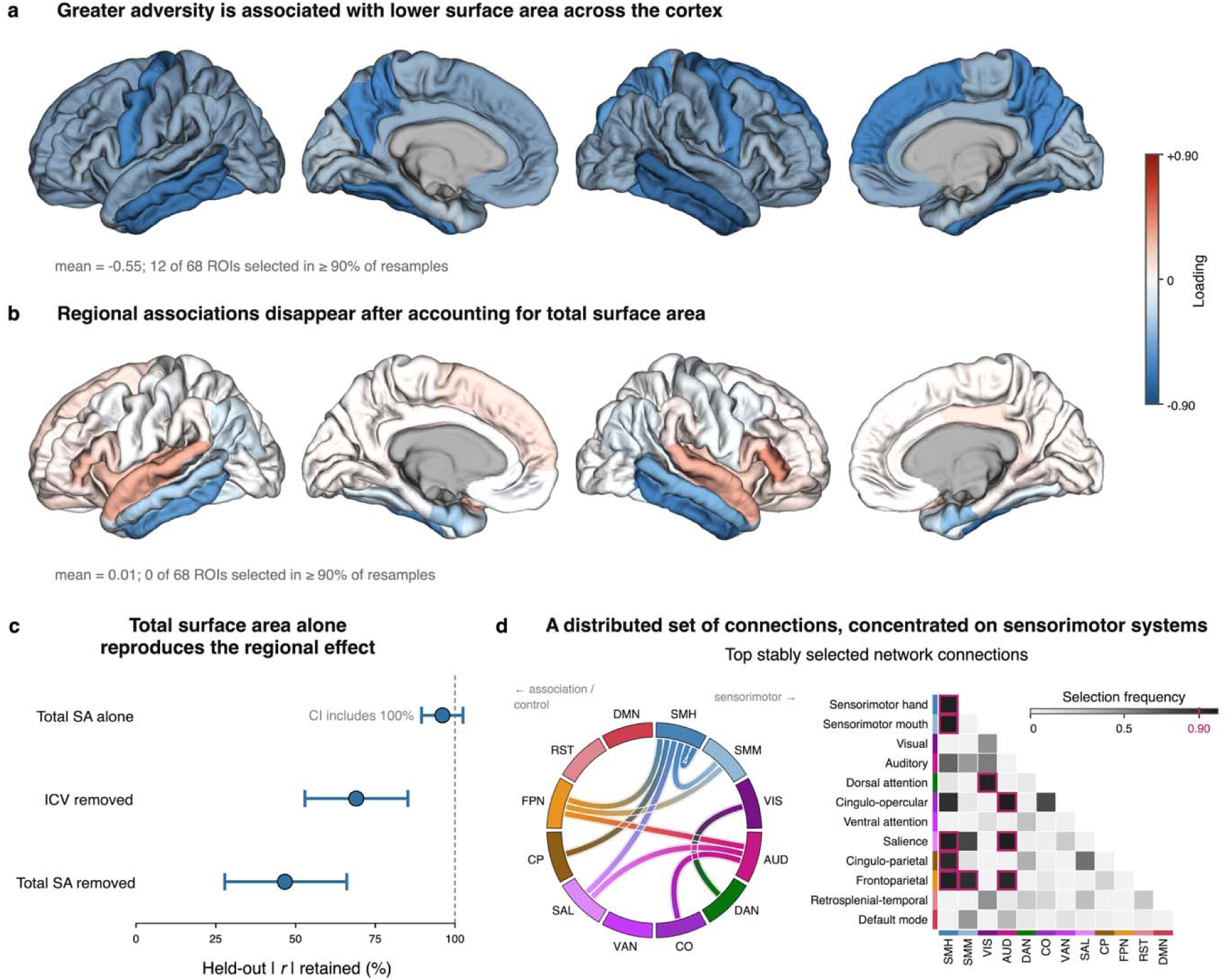
Socioeconomic Disadvantage Relates to Globally Reduced Cortical Surface Area and Sensorimotor-Weighted Connectivity. **(a)** Bootstrapped surface area loadings across 68 cortical regions. Loadings universally negative (mean = -0.55), indicating greater adversity is associated with lower surface area; 12 of 68 regions are stably selected (≥90% of resamples). **(b)** After removing total surface area, no regionally specific pattern survives (mean = 0.01; 0 of 68 regions stably selected), indicating a global rather than regional effect. **(c)** Association retained in held-out data after each adjustment. Total surface area alone reproduces the full effect (CI includes 100%), whereas removing intracranial volume or total surface area attenuates it, confirming a global surface area effect not attributable to head size. **(d)** Top stably selected resting-state connections (chord diagram and network-by-network matrix; ≥90% selection). Selected edges are distributed across the connectome but concentrated on sensorimotor systems.

The resting-state contribution localized to a reproducible set of connections rather than to a single global measure. Ten of 78 network pairs were stably selected in at least 90% of subsamples (13 at 75%; maximum 0.979), predominantly linking sensorimotor, auditory, and salience/cingulo-opercular systems (Fig. 3d): sensorimotor hand-sensorimotor mouth (0.979), auditory-cingulo-opercular (0.978), auditory-salience (0.971), within sensorimotor-hand (0.969), frontoparietal-sensorimotor hand (0.962), dorsal attention-visual (0.957), salience-sensorimotor hand (0.951), auditory-frontoparietal (0.929), cingulo-parietal-sensorimotor hand (0.911), and frontoparietal-sensorimotor mouth (0.901). Although this set of connections was reliably selected, no individual loading reached FDR significance (smallest *q*=.405), indicating that reliability resided in the distributed connectivity pattern rather than the magnitude of any single edge. Because ABCD’s preprocessing regresses out the global signal before computing network correlations, this pattern cannot be explained by a global increase or decrease in connectivity strength; instead, it reflects differences in the relative organization of connectivity across networks.

### Prediction of Year-6 Cognition, Psychopathology, and Physical Health

We next tested whether this socioeconomic disadvantage dimension predicted participants’ functioning six years later, adjusting for age at follow-up, sex, site, and, for cognition and psychopathology, the baseline value of the outcome. The adversity variate predicted lower fluid (*β*=-0.173 [-0.201, -0.144], *q*<.0001, ΔR^2^=0.035) and crystallized cognition (*β*=-0.095 [-0.118, -0.072], *q*<.0001, ΔR^2^=0.012), and greater waist-to-height ratio (*β*=0.214 [0.176, 0.253], *q*<.0001, ΔR^2^=0.045) and diastolic blood pressure (*β*=0.115 [0.079, 0.152], *q*<.0001, ΔR^2^=0.013). Systolic blood pressure (*β*=0.025, *q*=.20), total cholesterol (*β*=-0.026, *q*=.27), and glycated hemoglobin (*β*=0.058, *q*=.080) were not significant after correction (Fig. 4a). In addition, the variate was not associated with change from baseline in any CBCL scale (total *β*=-0.001, *q*=.93; internalizing *β*=0.017, *q*=.27; externalizing *β*=0.025, *q*=.15); without adjusting for baseline values, however, the socioeconomic disadvantage-related dimension was associated with year-6 internalizing, externalizing, and total problems (*β*s=0.12-0.16, all *p*<.001; Supplement Table S14). Thus, the brain-linked dimension forecasts poorer cognition and physical health, and tracks psychopathology levels already present at baseline, but does not predict symptom change across adolescence.

**Figure 4:**
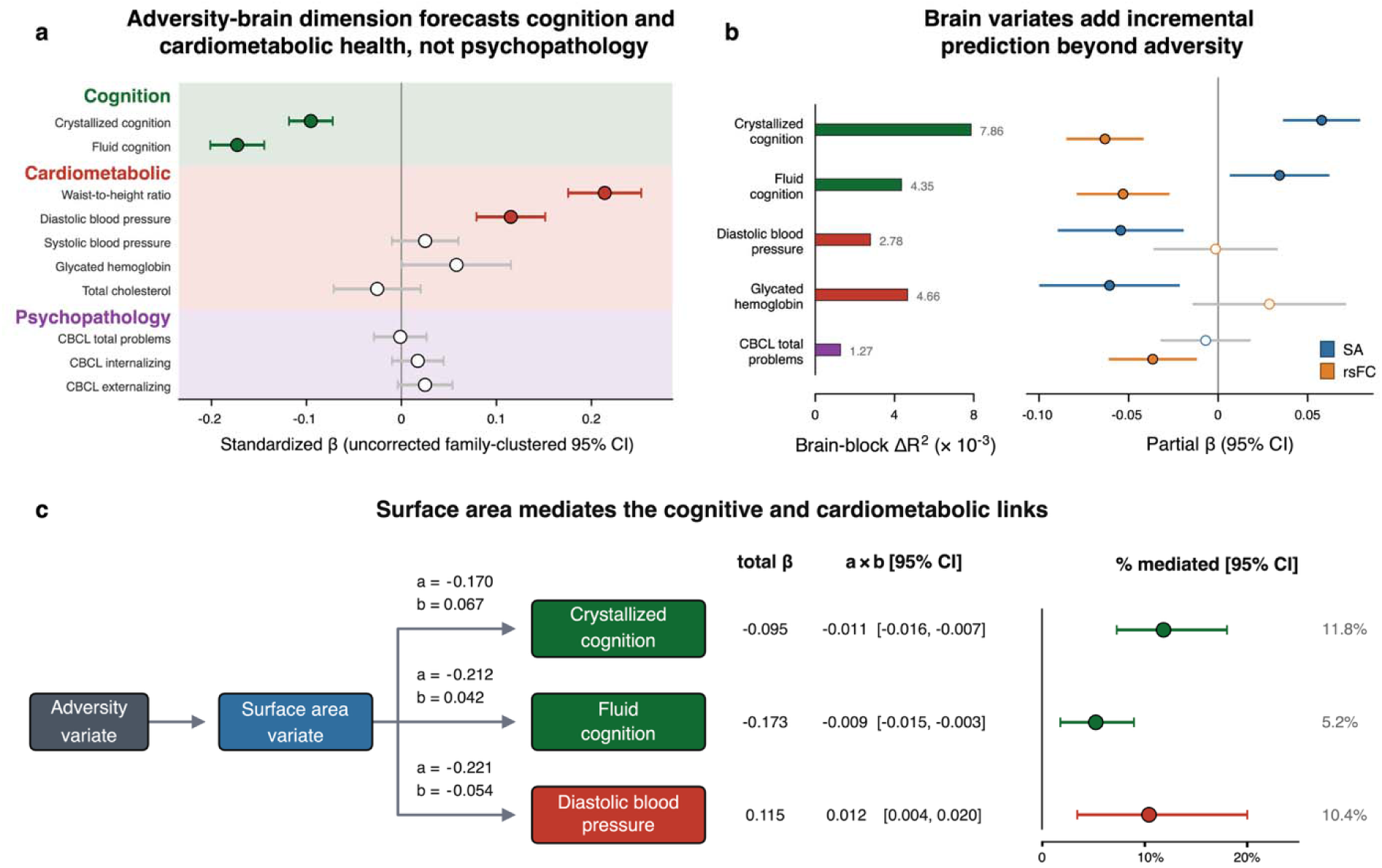
The Adversity-Brain Dimension Predicts Cognition and Cardiometabolic Health, but not Psychopathology. **(a)** Standardized associations between each individual’s position on the adversity-brain dimension and cognitive, cardiometabolic, and psychopathology outcomes (*β* ± uncorrected family-clustered 95% CI). The dimension predicts lower crystallized (-0.10) and fluid (-0.17) cognition and poorer cardiometabolic health (waist-to-height ratio +0.21; diastolic blood pressure +0.12), but not CBCL psychopathology (filled points = CI excludes 0). **(b)** Brain variates add predictive value beyond adversity: brain block ΔR^2^ (left) and partial *β* for the surface area and resting-state variates, adjusting for the adversity variate (right). **(c)** Cortical surface area partially mediates the dimension’s links to crystallized cognition (11.8% mediated), fluid cognition (5.2%), and diastolic blood pressure (10.4%); paths show a, b, total *β*, indirect effect (a x b), and percent mediated with 95% CIs.

To examine whether the brain variates carried information beyond adversity itself, we added both brain variates (surface area and resting-state connectivity) to a model containing the adversity variate alone. The two brain variates were only weakly and negatively correlated across individuals (r=-0.18), indicating they capture complementary rather than redundant information, consistent with two partially independent brain expressions of the same exposure dimension. The brain variates added modest but significant prediction for five outcomes (Fig. 4b): crystallized cognition (ΔR^2^=0.0079, *q*<.001), fluid cognition (ΔR^2^=0.0044, *q*<.001), glycated hemoglobin (ΔR^2^=0.0047, *q*=.024), diastolic blood pressure (ΔR^2^=0.0028, *q*=.024), and CBCL total problems (ΔR^2^=0.0013, *q*=.025), beyond adversity and covariates. For cognition, both variates contributed (crystallized: SA *β*=0.058, rsFC *β*=-0.063; fluid: SA *β*=0.034, rsFC *β*=-0.053), whereas the physical health and psychopathology increments were each carried by one modality (SA for glycated hemoglobin and diastolic blood pressure; rsFC for CBCL total). Thus, although the adversity dimension did not predict change in psychopathology, the rsFC variate accounted for a small additional proportion of variance in year-6 CBCL total problems beyond adversity. Systolic blood pressure, total cholesterol, waist-to-height ratio, and CBCL internalizing and externalizing showed no significant brain increment (all *q*>.05).

Finally, we tested whether the SA variate statistically mediates the adversity-outcome associations, bootstrapping the indirect effect over 5,000 family-clustered resamples. SA mediated a small share of the cognitive associations (Fig. 4c): 5.2% [1.7%, 8.9%] for fluid ability (indirect=-0.0090 [-0.0149, -0.0030], *q*=.005) and 11.8% [7.2%, 18.1%] for crystallized ability (indirect=-0.0113 [-0.0158, -0.0071], *q*=.002). A comparable indirect effect was present for diastolic blood pressure (10.4% [3.3%, 19.9%]; indirect=0.0120 [0.0040, 0.0204], *q*=.005). There was no indirect effect for waist-to-height ratio (*q*=.57) and, for glycated hemoglobin, the total effect was too near zero for a stable proportion mediated (95% CI extending to 91%). Therefore, mediation was modest and most consistent for cognition, with an additional significant effect for diastolic blood pressure.

### Robustness to Sensitivity Analyses

The socioeconomic disadvantage-related dimension and its principal brain associations were robust across alternative analytic specifications. Restricting the adversity battery to measures available at baseline, retaining one participant per family, and training on 15 sites while testing on six entirely held-out sites preserved the surface-area and rsFC associations, with held-out effects retaining 97-132% of the primary estimates (Supplementary Table S15; values above 100% reflect a larger estimate in the sensitivity sample). Removing site residualization from the adversity view produced highly similar environmental loadings and retained the same single dimension and modality specificity, although the SA and rsFC associations were modestly attenuated. The canonical solution was likewise preserved at the cross-validation optimal L1 penalty of 0.7, with the same dimensionality, environmental profile, and SA and rsFC specificity. Finally, the year-6 waist-to-height ratio association remained significant after adjustment for baseline waist-to-height ratio, although its magnitude was reduced by approximately 48%, indicating that a substantial portion of the adolescent association was already evident at ages 9-10 years (Supplementary Table S14).

## Discussion

Children who grow up in environments characterized by resource scarcity have been found to differ from their peers raised in more advantaged environments across a wide range of measures of brain structure and function. Because few studies of adversity have simultaneously considered a breadth of environmental exposures and neuroimaging modalities implicated in development, it is not clear whether these differences reflect many separable environmental influences or a smaller number of organizing dimensions, and whether they extend broadly across brain systems or are concentrated within a few regions. In this study we examined how the early environment is associated with neurodevelopment when both are measured comprehensively. We modeled 104 forms of early adversity spanning interpersonal, household, and neighborhood exposures against 7 structural and functional imaging modalities and tested whether the resulting dimensions replicated in held-out data. We obtained a single robust adversity-brain dimension defined, on the adversity side, primarily by socioeconomic and neighborhood deprivation, and on the brain side, by two imaging measures: lower cortical surface area and a distributed pattern of resting-state functional connectivity concentrated in sensorimotor systems. This dimension, measured at ages 9-10 years, predicted cognitive performance and cardiometabolic health six years later.

Neighborhood- and household-level measures contributed most strongly to this dimension; importantly, family conflict, neglect, and parenting behavior were not reliably selected. The highest loading measures assessed child opportunity, social mobility, neighborhood deprivation, and social vulnerability, followed by household income and caregiver education, pointing to socioeconomic resources and the broader opportunity structure surrounding a child as the principal environmental features defining this axis. Given that all exposures were modeled simultaneously, this pattern suggests that broader structural and neighborhood-level conditions are associated more strongly with neurodevelopmental variation than are more proximal family-level exposures. The predominance of measures indexing socioeconomic resources and opportunity further aligns the brain-linked environmental axis most closely with the deprivation dimension of the threat-deprivation framework^15,28^. Notably, in one of the earliest large-scale studies of the relation between socioeconomic variation and the developing brain, Noble and colleagues (2015)^10^ linked family income and parental education to cortical SA. We found here that a similar SA signature was the most robust structural correlate from a substantially broader model of environmental experience and neurodevelopment. This association in the brain was largely global; total cortical SA alone reproduced 96% of the out-of-sample regional effect, and no individual region survived selection after total SA was residualized from each parcel. Thus, socioeconomic disadvantage appears to be associated more consistently with variation in overall cortical surface area than with regionally specific differences in cortical morphology.

Resting-state connectivity revealed a complementary functional signature of socioeconomic disadvantage centered on sensorimotor systems. Marek and colleagues^29^ recently examined several hundred environmental, behavioral, and demographic measures in the same cohort and identified socioeconomic status as the strongest environmental correlate of rsFC, with associations concentrated in sensory and motor systems rather than in higher-order association cortex. The connections that were selected most consistently in the present study similarly involved sensorimotor, attention, and salience systems. Our multimodal analysis extends these findings by demonstrating that when seven imaging modalities are evaluated jointly and are required to replicate in held-out families, the socioeconomic gradient is reproducibly expressed in SA and rsFC, but not in cortical thickness, white-matter fractional anisotropy, or activation during any of the three fMRI tasks. This selectivity should not be interpreted to mean that socioeconomic disadvantage is unrelated to these other neural measures; instead, it indicates that their linear multivariate associations were not sufficiently strong or stable to contribute reliably when examined alongside the other modalities. The contrast between SA and cortical thickness is particularly notable, suggesting that associations between socioeconomic conditions and cortical morphology during late childhood are reflected more consistently in the extent of the cortical sheet than in its thickness. Moreover, the weak correlation between the SA and connectivity variates indicates that these structural and functional signatures capture largely distinct aspects of neurodevelopmental variation associated with socioeconomic disadvantage.

The socioeconomic disadvantage-related dimension identified in late childhood also predicted cognitive and physical health six years later. Greater expression of this dimension at ages 9-10 years predicted poorer cognitive performance in adolescence, particularly crystallized cognition, as well as greater waist-to-height ratio and higher diastolic blood pressure. The cognitive findings are consistent with socioeconomic differences in the cumulative acquisition of language, knowledge, and educational skills across childhood^30–32^. Similarly, the cardiometabolic findings echo longstanding evidence linking adverse childhood environments to later cardiometabolic disease risk^33,34^ and indicate that the socioeconomic disadvantage-related dimension identified here is associated with emerging differences in physical health by late adolescence^35^. SA statistically mediated a modest proportion of the prospective associations with cognition and diastolic blood pressure, suggesting that adversity-related variation in cortical SA represents a potential neurodevelopmental pathway linking early socioeconomic conditions to later cognitive and physical health. The adversity-brain dimension did not, however, predict changes in parent-reported psychopathology after accounting for baseline symptoms, despite associations with follow-up symptom levels in models that did not adjust for baseline symptoms. Thus, the dimension appears to track individual differences in mental health that were already present by late childhood, but not the subsequent progression of symptoms over the period examined.

Our findings also raise questions about the utility of current dimensional taxonomies for understanding the neurodevelopmental consequences of adversity. Within this multivariate design, we did not recover additional reproducible brain-linked dimensions corresponding to threat, unpredictability, or proximal family experiences. Frameworks distinguishing threat, deprivation, and unpredictability were developed, in part, to explain why different adverse experiences may have different consequences for behavioral and psychobiological development^36–38^. While our results support the formulation that adverse experiences are not interchangeable, given the selective association of deprivation with neurodevelopment, they provide little support for the multidimensional organization proposed by these frameworks. Across the full environmental battery, deprivation was the only dimension with a reproducible multivariate association with the brain; no additional dimensions emerged corresponding to threat, unpredictability, or more proximal family experiences. When they are examined separately, associations that are identified between theoretically defined adversity dimensions and different neural outcomes do not establish that these dimensions represent independent or separable patterns of adversity-related neurodevelopment. Because environmental exposures and neural phenotypes are highly correlated, such specificity is better tested by modeling their covariance simultaneously. Using this more stringent test in the present study, we did not find the multidimensional structure that is predicted by current frameworks. Therefore, adversity neuroscience may be better served by deriving dimensions from reproducible, taxonomy-agnostic relations between environmental variation and neurodevelopment than by imposing classifications of adverse experience *a priori* and assuming that they map onto distinct features of the developing brain.

We should note four limitations of this study. First, because the data we analyzed here are observational, the associations we identified should not be interpreted to mean that socioeconomic disadvantage *causes* differences in neurodevelopment or later health. Second, substantial correlations among environmental measures, reflecting the well-documented co-occurrence of adversities and socioeconomic risks^13,14^, may limit inferences about the contribution of any individual exposure. Although the multivariate weights identify measures that characterize the socioeconomic disadvantage-related dimension most consistently, they cannot determine which correlated environmental features independently account for the association of this dimension with the brain. Third, we did not examine explicit genetic factors; socioeconomic circumstances covary with heritable parental characteristics that may also be related to offspring neurodevelopment, and the limited within-family variability in socioeconomic and neighborhood exposures prevents sibling comparisons from separating these influences in the present data. Finally, the analytic sample was more likely to be white and socioeconomically advantaged than the full ABCD cohort, consistent with previously reported selection patterns in complete-case neuroimaging samples in this cohort^27^. This sampling pattern may have restricted environmental variation, potentially attenuating associations and limiting generalizability to children who were experiencing more severe socioeconomic disadvantage.

Despite these limitations, this study is important in demonstrating that the relation between early adversity and neurodevelopment is substantially lower-dimensional than the breadth of reported adversity-brain associations might imply. Across a wide range of environmental experiences and seven imaging modalities, a single reproducible dimension emerged, organized around socioeconomic and neighborhood deprivation and characterized by lower cortical SA and altered rsFC centered on sensorimotor systems. Further, prospective associations with cognitive and cardiometabolic outcomes indicate that this dimension captures developmental differences that remain consequential into late adolescence. Rather than a diffuse set of exposure- and modality-specific associations, our findings indicate that when the early environment and the developing brain are modeled as an integrated system, socioeconomic deprivation is identified as the principal environmental dimension organizing reproducible variation in brain structure and function.

## Methods

### Participants and Study Design

Participants were drawn from the Adolescent Brain Cognitive Development (ABCD) Study^39^ (Release 7.0; https://abcdstudy.org/), a longitudinal cohort of 11,860 youth enrolled at ages 9-10 years across 21 sites in the United States. The cohort excluded youth with sensory, neurological, medical, or intellectual limitations that would preclude completion of the study protocol, and any MRI contraindications. At the baseline visit, participants completed a broad assessment of the early environment together with structural, diffusion-weighted, and functional MRI; at the 6-year follow-up, they completed measures of cognition, psychopathology, and physical health. The early-environment assessment was composed of 104 measures spanning interpersonal, household, and neighborhood levels. All data were drawn from measures administered at the baseline visit except the Life Events Scale (year-1 follow-up visit) and the neglect measure (year-3 follow-up visit), each taken from its earliest available assessment; baseline-only sensitivity analyses are reported in the Supplement. Neuroimaging yielded seven feature sets across the structural, diffusion, and functional modalities (see Measures). The primary analytic sample was composed of the 5,976 participants (from 5,303 families) with complete neuroimaging data across all seven modalities. Follow-up sample sizes varied by measure and were nested within this sample: 4,846-4,849 for psychopathology (Child Behavior Checklist^40^), 3,718-3,851 for cognition (NIH Toolbox^41^), and 1,565-3,101 for physical health (waist-to-height ratio, blood pressure, glycated hemoglobin, and total cholesterol). Demographic characteristics of the sample are presented in Table 1.

### Measures

#### Adversity and the Early Environment

We adapted the adversity battery from Brieant et al.’s (2023)^20^ list of 60 baseline measures spanning experiential and environmental exposures, and extended it in two ways. First, we added two instruments assessing exposures not captured at baseline, including the Life Events Scale^42^ (year-1 follow-up) and the Multidimensional Neglectful Behavior Scale^43^ (year-3 follow-up). Second, we added a curated set of linked external data (LED) measures tied to each participant’s primary residential address at baseline, newly available as part of ABCD Release 7.0. The resultant battery was composed of 104 features, providing a comprehensive representation of the early environment across interpersonal, household, and neighborhood levels.

Features were drawn from the Life Events Scale^42^ (parent report; 25 items); the computerized Kiddie Schedule for Affective Disorders and Schizophrenia for DSM-5^44^ (KSADS) traumatic-events screener (18 items); the conflict subscale of the Family Environment Scale^45^, administered to both caregiver and youth (18 items, 9 per informant); the Multidimensional Neglectful Behavior Scale^43^ (youth report; 8 items); the DSM-oriented subscales of the Adult Self-Report^40^, indexing caregiver ADHD, antisocial, anxiety, avoidant, depressive, and somatic symptoms (6 scales); the Family History Assessment^46^, indexing maternal and paternal alcohol and drug problems (4 items); the Neighborhood Safety and Crime Scale^47^, caregiver and youth report (2 scales); the acceptance subscale of the Children’s Report of Parental Behavior Inventory^48^ for each of two caregivers (2 scales); and the Parental Monitoring Scale^49^ (1 scale). Nine features indexed household socioeconomic status: household income, caregiver education, partner education, and six indicators of past-year financial hardship (inability to afford food, telephone service, rent or mortgage, utilities, or medical care, and eviction from the home). A further 11 LED^50^ features characterized the neighborhood environment: area deprivation, child opportunity, social vulnerability, social mobility, nitrogen dioxide and fine particulate matter concentrations, lead exposure risk, noise, tree canopy, alcohol outlet density, and crime.

We selected LED measures indexing socioeconomic, environmental hazard, and neighborhood resource domains not captured by item-level self-report and not included by Brieant et al. (2023)^20^. Two candidate measures, walkability and ozone, were excluded because their *a priori* adversity orientation was not supported; neither tracked household socioeconomic disadvantage in the expected direction (*r*=-0.13 and -0.03), as less walkable areas tended to be car-dependent, higher-income suburbs and ground-level ozone is higher in less urban, more affluent areas. Every feature was coded such that higher values indicate greater adversity, which required reverse-scoring the protective constructs (caregiver warmth, parental monitoring, neighborhood safety, child opportunity, social mobility, and tree canopy). ABCD ‘decline-to-answer,’ ‘don’t-know,’ and other implausible sentinel codes were recoded as missing; the remaining missing adversity values (3.2% of adversity view cells, no feature exceeding 16.3%) were imputed with the training set median, and features were standardized using the training set mean and standard deviation, applied unchanged to the test set. The seven neuroimaging views contained no missing data (the analytic sample was their complete-case intersection) and outcomes were never imputed. Supplementary Table S1 lists all 104 features with the source instrument, ABCD variable name, coding direction, and stability-selection frequency.

#### Neuroimaging Measures

##### Acquisition and Processing

Structural, diffusion, and functional MRI were acquired at the baseline visit on 3T Siemens, Philips, and GE scanners using a harmonized multiband protocol; full acquisition parameters and the scan protocol are provided in Casey et al. (2018)^51^ and Supplementary Table S2. All measures were taken from the ABCD tabulated Release 7.0 derivatives produced by the centralized DAIRC pipeline^52^, which applies corrections for gradient nonlinearity, susceptibility-induced distortion, and head motion. Cortical surfaces were reconstructed and subcortical and cerebellar structures segmented with FreeSurfer 7.1.1^53^, with the cortex parcellated by the Desikan-Killiany atlas^54^; white matter tracts were labeled with AtlasTrack^55^; and resting-state functional connectivity was computed as Fisher *z*-transformed correlations among the Gordon parcellation networks^56^, after nuisance regression of motion, white matter, cerebrospinal fluid, and the whole-brain (global) signal. Each modality was filtered using its own DAIRC quality-control inclusion criteria, and participants failing the threshold for any modality were excluded.

##### Features

Seven neuroimaging views were retained for modeling. Cortical thickness (CT) and cortical surface area (SA) were composed of the 68 Desikan-Killiany parcels (34 per hemisphere). Fractional anisotropy (FA) was composed of 37 features, including 17 white matter tracts per hemisphere (34), the corpus callosum, forceps major, and forceps minor. Resting-state functional connectivity (rsFC) contributed 78 features including 12 within-network correlations and 66 between-network correlations among the 12 Gordon networks. Each of the three task fMRI views provided 84 features from a single canonical contrast – anticipation of a large reward versus neutral (MID), correct stop versus correct go (SST), and emotional versus neutral face (ENB) – including the 68 Desikan cortical parcels and 16 left/right gray matter structures (accumbens, amygdala, caudate, hippocampus, pallidum, putamen, thalamus, and cerebellar cortex); white matter, CSF, ventricular, ventral diencephalon, and brainstem parcels were excluded. Total cortical SA (the sum of the 68 regional SA features) was retained separately for the global-versus-regional sensitivity analysis described below.

#### Covariates

We regressed age, sex, and study site from every feature view (the adversity view and all seven neuroimaging views), estimating each residualization model in the training set and applying it unchanged to the test set. Study site was residualized from the adversity view as well as the neuroimaging views to prevent between-site differences in environmental composition and imaging acquisition from driving cross-view associations. Race and ethnicity were not included as covariates because they are closely intertwined with the socioeconomic and neighborhood conditions under study and were not conceptualized as nuisance sources of variation to be removed from the environmental dimension. For the five views in which head motion is a plausible source of artifact and mean framewise displacement (FD) was available in the tabulated release data (rsFC, the three task views, and FA), we additionally regressed out that modality’s mean FD. Residuals were then standardized and each view was divided by the square root of its number of features, so that no view influenced the canonical solution simply by containing more features. This full procedure was repeated within each sensitivity analysis, using only the training data from that analysis.

#### Outcome Measures

We examined outcomes across three domains at the Year-6 follow-up visit. First, we assessed mental health using the parent-reported Child Behavior Checklist (CBCL)^40^, a well-validated measure of emotional and behavioral problems in youth aged 6-18 years; three broadband raw scores (Total Problems, Internalizing, and Externalizing) served as outcomes. Second, we indexed cognitive ability using the NIH Toolbox Cognition Battery^41,57^, a standardized, computerized battery normed to a nationally representative US sample. The fluid composite aggregates five tasks assessing processing speed, attention and inhibitory control, working memory, cognitive flexibility, and episodic memory; the crystallized composite reflects accumulated verbal knowledge from picture vocabulary and oral reading recognition subtests. Both are expressed as uncorrected standard scores. Third, we assessed cardiometabolic health, indexed by five markers: waist-to-height ratio, systolic and diastolic blood pressure, total cholesterol, and glycated hemoglobin (HbA1c). Because early adversity is a well-documented risk factor for cardiometabolic disease and premature mortality in adulthood^33,34^, we examined these markers to capture environment-linked physiological risk that may become embedded during adolescence, years before clinical disease emerges.

For psychopathology and cognition, which were also assessed at baseline, we included the baseline value of each measure as a covariate in the corresponding year-6 model; thus, those estimates reflect prospective prediction of change rather than prediction of raw values. Most cardiometabolic measures (blood pressure, cholesterol, and HbA1c) were first assessed after baseline, so for comparability across the cardiometabolic block we did not include a baseline covariate for any of these outcomes. To maximize power, each model included all participants with the relevant year-6 outcome; therefore, sample sizes vary across outcomes (see Participants) and are nested within the primary sample.

### Statistical Analyses

#### Sparse Generalized Canonical Correlation Analysis

Rather than testing each imaging feature against each adversity measure separately, we used sparse generalized canonical correlation analysis (SGCCA)^58,59^, a data-driven multivariate approach, to characterize how the early environment, measured broadly, maps onto brain structure and function. SGCCA operates on the eight views (the adversity view and seven imaging views) and assigns each feature a weight, choosing weights that maximize the association between the resulting view variates; thus, each component expresses a pattern of adversity features that covaries with a pattern of brain features. Following Lett et al. (2025)^60^, we used the factorial scheme g(x)=x^2^, a fully connected design (C=1-I), and successive deflation to extract components, with an L1 penalty^61^ shrinking most weights to zero and retaining a parsimonious contributing set.

To guard against overfitting and respect the embedded sibling structure of the ABCD cohort, we randomly assigned families to a 70:30 training-test split (training: 4,186 youth in 3,712 families; test: 1,790 in 1,591 families) so that no siblings were included in both samples. The training and testing sets were comparable on sociodemographic characteristics and the primary study variables (Table 1). All model fitting, penalty tuning, stability selection, and bootstrapping was performed in the training set. We selected the L1 penalty by 5-fold, family-grouped cross-validation in the training set; although the cross-validation optimum was L1=0.7, the canonical criterion was flat across the 0.4-0.7 range (Δ=0.0016, *SE*=0.0056; within resampling noise), so we fixed L1=0.4 for greater sparsity and interpretability. We extracted five candidate components to allow for multiple adversity-brain dimensions and assessed the significance of each by permutation (10,000 draws), using as the test statistic the average variance explained across the seven adversity-brain pairs (minimum attainable *p*=.0001). In the training set, the adversity view was permuted and the model refit; in the test set the fixed training-derived weights were applied and the adversity rows permuted, so held-out *p*-values reflect the permutation null rather than any re-estimated model. A component was retained only if it exceeded its permutation null in both the training and held-out sets. Because the sign of a variate is arbitrary, associations are reported as |*r*| and each variate is then contextualized by its dominant loadings. Because the L1 penalty biases individual variate scores, all subject-level analyses (the held-out scatterplots in Fig. 1d and the year-6 outcome models) used scores from a companion model refit without sparsity (L1=1) on the same views and training/test split; the sparse model was retained for feature selection and loading interpretation.

To identify reliable features, we refit the model in 10,000 subsamples of 70% of training families and recorded how often each feature received a non-zero weight; following Lett et al. (2025)^60^, features selected in ≥90% of subsamples are considered stably selected. To characterize each variate, we computed structural loadings (the correlation of each feature with its own view’s variate) over 10,000 family bootstrap draws, with 95% CIs and Benjamini-Hochberg (BH) FDR correction. For views in which all features loaded in the same direction, the variate is essentially a weighted average of that view, and virtually every loading is statistically significant; because significance is then uninformative for distinguishing among features, we instead interpreted features by the relative magnitude of their loadings and their stability-selection frequency.

#### Year-6 Outcome Models

Each year-6 outcome was regressed on each significant adversity variate (each participant’s score on the SGCCA-derived adversity dimension) and on the associated brain variate using ordinary least squares with family-clustered robust standard errors and covarying age at the year-6 assessment, sex, site, and, for psychopathology and cognition, the baseline value of the outcome. Motion was not included, as the brain variates were already motion residualized. Predictors and outcomes were z-scored within each model, so coefficients are standardized betas, and BH-FDR was applied across outcomes within each predictor and domain. To test whether the brain variates carried information beyond adversity, we compared a model containing the adversity variate alone with one that additionally included both brain variates (surface area and resting-state connectivity), evaluating the change in R^2^ for the brain block and the standardized partial coefficient of each brain variate. We then tested whether each brain variate statistically mediated the adversity-outcome association (indirect effect over 5,000 family bootstrap draws).

#### Sensitivity Analyses

We repeated the full analytic pipeline under several alternative specifications, each designed to test the robustness of the primary findings. First, we restricted the adversity battery to measures collected at baseline to determine whether the identified dimension depended on the two instruments administered at later assessments. Second, we repeated the analysis after randomly retaining one participant per family to assess whether the canonical solution depended on the inclusion of related participants. Third, we trained the model using participants from 15 study sites and evaluated it in participants from 6 entirely held-out sites, providing a more stringent test of generalization across sites and scanners. For each analysis, we quantified effect retention as the held-out |*r*| from the sensitivity analysis divided by the corresponding primary held-out |*r*| and expressed as a percentage. Values near 100% indicate preservation of the primary effect, whereas values above 100% reflect a larger estimate in the sensitivity sample rather than evidence of a stronger underlying association. Additional analyses reported in the Supplement included a within-family decomposition to characterize within- and between-family variation, adjustment for intracranial volume to assess the contribution of overall head size, a threat-versus-deprivation comparison to test for a separable threat-related dimension, and a cross-modal convergence analysis to assess whether the surface-area and connectivity signatures reflected a common environmental gradient.

## Data Availability

Data from the ABCD Study, including derivatives processed as part of ABCC, are available through the NIH Brain Development Cohorts (NBDC) Data Hub (https://www.nbdc-datahub.org/) and require a Data Use Agreement for access. Analyses used ABCD Release 7.0.

## Code Availability

All code used for the analyses described in this manuscript are available in an online repository (https://github.com/cantonacci/ABCD_environment-brain). Analyses were implemented in Python 3.12.1. Sparse generalized canonical correlation analysis was fit with an in-house Python implementation of the RGCCA/SGCCA framework (Tenenhaus & Tenenhaus, 2017)^59^, which is provided with the analysis code. The pipeline used NumPy 2.2.6, pandas 2.3.2 and SciPy 1.16.3 for data handling and numerical computation; statsmodels 0.14.6 for the outcome regression models with family-clustered standard errors and for FDR correction; and scikit-learn 1.7.2 for family-grouped cross-validation. For neuroimaging analyses, it used nibabel 5.4.2 and neuromaps 0.0.7. Figures were made with Matplotlib 3.11.1, and cortical surfaces were rendered with surfplot 0.2.0 (BrainSpace 0.2.1, VTK 9.6.2).

## Funding Statement

This work was supported by the National Institute of Mental Health (F31MH146295 to CA; R37MH101495 and R01MH140425 to IHG; F32MH135657 to JPU), the National Science Foundation (Graduate Research Fellowship Program to EG), and the Stanford Interdisciplinary Graduate Fellowship (to CA).

## Acknowledgements

We thank the participating families and ABCD research staff for their time and effort.

## Author Contributions Statement

Conceptualization (CA, JPU, IHG); Data curation (CA, KK); Methodology (CA, KK, EG, JPU, IHG); Formal analysis (CA, KK); Validation (CA, KK); Writing – original draft (CA, EG, IHG); Writing – review and editing (CA, KK, EG, JPU, IHG); Funding acquisition (CA, EG, JPU, IHG).

## Competing Interests Statement

The authors declare no competing interests.

## Notes

### Competing Interest Statement

The authors have declared no competing interest.

