## Supplementary Material for "A Single, Reproducible Dimension Organizes Associations Between the Early Environment and the Developing Brain"

**Contents**

**Supplementary Methods**

- Adversity and Environmental Battery
- Neuroimaging Features
- Covariate Residualization and Preprocessing
- SGCCA Specification, Penalty Selection, and Family-Aware Design
- Permutation Testing, Stability Selection, and Bootstrap Loadings
- Non-Sparse Companion Variates
- Supplementary Brain-Side Analyses
- Year-6 Outcome Models and Mediation
- Sensitivity Analyses

**Supplementary Results**

- Representativeness of the Analytic Sample
- Component Structure and Modality Specificity
- Environmental Feature Selection
- Global Versus Regional Cortical Surface Area
- Resting-State Connectivity
- Cross-Modal Convergence
- Threat Versus Deprivation
- Within-Family Decomposition
- Year-6 Outcomes: Baseline Adjustment, Brain Increments, and Mediation
- Baseline-Only Battery, One Participant per Family, and Held-Out Sites
- Site Residualization of the Adversity View
- Penalty Sensitivity (L1 = 0.7)

**Supplementary Tables**

- Table S1. The 104-Feature Adversity and Environmental Battery: Source, Coding, Selection, and Loadings
- Table S2. Harmonized MRI Acquisition Parameters by Scanner Platform
- Table S3. Neuroimaging Feature Sets and Native-Unit Descriptive Statistics
- Table S4. Penalty Selection by Five-Fold Family-Grouped Cross-Validation
- Table S5. Component 1 by View: Sparsity, Stability, and Training and Held-Out Associations
- Table S6. Analytic Sample Versus Excluded Baseline Participants
- Table S7. Global Versus Regional Decomposition of the Surface-Area Association
- Table S8. Regional Surface-Area Loadings and Selection Before and After Total Surface-Area Adjustment
- Table S9. Resting-State Network-Pair Loadings and Stability Selection (All 78 Features)
- Table S10. Cross-Modal Convergence of Single-Modality Adversity and Brain Axes
- Table S11. Threat Versus Deprivation Composites and Their Brain Associations
- Table S12. Within-Family Decomposition in Sibling Families
- Table S13. Year-6 Outcomes: Adversity Associations, Brain Increments, and Surface-Area Mediation
- Table S14. Baseline Adjustment of Year-6 Psychopathology and Waist-to-Height Ratio Associations
- Table S15. Baseline-Only, One-Per-Family, and Held-Out-Site Sensitivity Analyses
- Table S16. Primary Model Versus Site-Unresidualized Adversity View and L1 = 0.7 Penalty

**Supplementary Figures**

- Figure S1. Resting-State Network-Pair Loadings and the Global Surface-Area Association

**Supplementary References**

### Supplementary Methods

#### Adversity and Environmental Battery

The adversity view extended the 60 baseline measures of Brieant et al.^1^ by adding two instruments that assess exposures not captured at the baseline visit and 11 linked external data (LED) measures describing each participant’s residential neighborhood. Table S1 lists all 104 features. For presentation they are grouped into the four domains used in the main-text figures: interpersonal and life events (72 features: 25 Life Events Scale items^2^, 18 items of the KSADS traumatic-events screener^3^, 18 Family Environment Scale conflict items^4^ reported separately by caregiver and youth, 8 items of the Multidimensional Neglectful Behavior Scale^5^, 2 Children’s Report of Parental Behavior Inventory (CRPBI) acceptance scales^6^, and the Parental Monitoring Scale^7^); caregiver psychopathology and substance use (10 features: 6 DSM-oriented Adult Self-Report scales^8^ and 4 Family History Assessment indicators^9^); household socioeconomic status (9 features); and neighborhood and built environment (13 features: 2 Neighborhood Safety and Crime Scale scores^10^ and 11 LED measures^11^). The domain grouping is descriptive and played no role in model fitting, which treated the 104 features as a single view.

All features were taken from the ABCD Study^12^ baseline visit except the Life Events Scale, which was first administered at the year-1 follow-up, and the neglect scale, first administered at the year-3 follow-up; each was taken from its earliest assessment. LED measures were linked to the primary residential address recorded at baseline. Item-level instruments entered as individual binary items (event occurred or endorsed, conflict statement endorsed). Family Environment Scale items were used in the release’s harmonized coding, in which reverse-worded statements are already keyed in the conflict direction. Multi-item scales entered as composites: the mean of the five CRPBI acceptance items for each caregiver, the mean of the five parental-monitoring items, and the mean of the three caregiver-reported neighborhood-safety items (youth report is a single item). Family-history features indicated any reported alcohol or drug problem for the biological mother or father, and caregiver psychopathology entered as raw DSM-oriented scale sums. Every feature was coded so that higher values indicate greater adversity; the protective constructs (caregiver acceptance, parental monitoring, neighborhood safety, supervision and educational support, household income and education, child opportunity, social mobility, and tree canopy) were reverse-scored. The coding direction of each feature is given in Table S1.

ABCD ‘decline to answer,’ ‘don’t know,’ and other sentinel codes were recoded as missing. For continuous LED counts, a sentinel value was recoded only when it was implausible for that feature, so that a genuine count equal to a sentinel number was retained. Binary items endorsed by fewer than six participants were removed before modeling. Two candidate LED measures, walkability and ozone, were excluded because their *a priori* adversity orientation was not supported by their association with a household socioeconomic core that contained no LED features (main text, Methods). After cleaning, 3.2% of adversity cells (19,891 of 621,504) were missing, spread over 100 of the 104 features and 2,412 participants; the most incomplete feature was partner education (16.3%). Missing values were imputed with the training set median of each feature, and features were standardized with the training set mean and standard deviation; both were applied unchanged to the held-out set. The percentage of missing values for each feature is listed in Table S1. Imaging features and outcomes were never imputed.

#### Neuroimaging Features

MRI data were acquired at the baseline visit on Siemens, Philips, and GE 3T scanners using the harmonized ABCD protocol^13^; Table S2 lists the acquisition parameters for each platform. All imaging features were taken from the tabulated Release 7.0 derivatives of the centralized DAIRC pipeline^14^. For each modality we applied ABCD’s recommended quality-control inclusion flag (T1-weighted structural data for cortical thickness and surface area, diffusion data for FA, and the resting-state and each task run for the functional views). Before intersection, usable data were available for 11,554 participants for the structural views, 10,404 for FA, 9,554 for rsFC, 9,256 for the monetary incentive delay (MID) task, 8,099 for the stop-signal task (SST), and 7,819 for the emotional n-back (ENB) task. The analytic sample is the complete-case intersection of all views (N = 5,976).

Table S3 summarizes the seven feature sets. Cortical thickness (CT) and surface area (SA) comprised the 68 Desikan-Killiany parcels^15,16^. FA comprised 37 AtlasTrack tract values^17^ (17 tracts in each hemisphere, the corpus callosum, and the forceps major and minor). rsFC comprised the 78 unique within- and between-network mean correlations among the 12 Gordon communities^18^ (auditory, cingulo-opercular, cingulo-parietal, default mode, dorsal attention, frontoparietal, retrosplenial-temporal, salience, sensorimotor hand, sensorimotor mouth, ventral attention, and visual). Each network pair entered once, and correlations involving parcels not assigned to a Gordon community were excluded. Each task view comprised the 84 gray matter parcel estimates of one contrast (68 cortical and 16 subcortical and cerebellar structures). Two global structural measures were retained outside the model for the global-versus-regional analyses. Total cortical SA was the sum of the 68 regional SA features (mean = 190,289 mm^2^, SD = 17,661; coefficient of variation = 9.3%). Intracranial volume (ICV) was FreeSurfer’s estimated total intracranial volume, which correlated *r* = 0.767 with total SA.

#### Covariate Residualization and Preprocessing

Within the training set, each feature of every view was regressed on age at baseline, sex, and study site (21 sites, dummy coded) by ordinary least squares, and the training coefficients were applied unchanged to the held-out set. For the rsFC, FA, and three task views, the covariate set also included that modality’s mean framewise displacement (FD), or, for FA, the diffusion head motion estimate. The structural views received no motion covariate. Residuals were standardized with training set means and standard deviations, and each view was then divided by the square root of its number of features, so that views with more features did not dominate the joint criterion. Every sensitivity analysis repeated this procedure from the start, using only its own training rows.

Study site was residualized from the adversity view as well as the neuroimaging views to prevent between-site differences in environmental composition and imaging acquisition from driving cross-view associations. A consequence is that the adversity variate indexes differences in environment among children recruited at the same site (the site-residualization sensitivity analysis below tests robustness to this decision). Race and ethnicity were not included as covariates because they are closely intertwined with the socioeconomic and neighborhood conditions under study and were not conceptualized as nuisance sources of variation to be removed from the environmental dimension.

#### SGCCA Specification, Penalty Selection, and Family-Aware Design

Sparse generalized canonical correlation analysis (SGCCA)^19,20^ was applied to the eight views (the adversity view and the seven imaging views). For component 1, SGCCA finds one weight vector per view that maximizes the sum, over all 28 pairs of views, of the squared covariance between the two views’ variates (the factorial scheme, g(x) = x^2^, with a fully connected design, C = 1 - I)^21^. Each weight vector has unit L2 norm and an L1 norm no larger than s√p, where p is the number of features in the view and s is the penalty parameter; s = 1 imposes no sparsity, and smaller values shrink more weights to exactly zero^22^. The solution was obtained by block coordinate ascent with soft threshold projection (convergence tolerance 10^-8^ on the relative change in the criterion, maximum 1,000 iterations). Later components were extracted by successive deflation, in which each view was regressed on its own component score before the next component was fitted. Five candidate components were extracted so that additional adversity-brain dimensions, such as separable threat- or family-related dimensions, could emerge if present. Each was retained only if it replicated in both the training and held-out sets. SGCCA estimates linear multivariate associations; it does not model nonlinear effects.

Families (not individuals) were randomly assigned to a 70:30 training/held-out split, so that no siblings were divided across the two sets (training: 4,186 youth in 3,712 families; held-out: 1,790 youth in 1,591 families). The penalty was selected by 5-fold cross-validation within the training set, with folds grouped by family, over the grid s ∈ {0.1, 0.2, 0.3, 0.4, 0.5, 0.7}. In each fold the model was fitted on 80% of the training families, and the criterion was the mean absolute correlation between the adversity variate and each of the seven brain variates in the remaining 20% (Table S4). The criterion peaked at s = 0.7 but was flat between 0.4 and 0.7: the paired fold difference was 0.0016 (*SE* = 0.0056), well within resampling noise. We therefore fixed s = 0.4, which yields a sparser solution in which stability selection remains informative. At s = 0.7, 103 of the 104 adversity features receive non-zero weights (see Penalty Sensitivity below). All model fitting, tuning, stability selection, and bootstrapping used the training set only.

#### Permutation Testing, Stability Selection, and Bootstrap Loadings

The test statistic for each component was the mean squared correlation between the adversity variate and each of the seven brain variates, that is, the mean variance explained across the seven adversity-brain pairs. The 21 brain-brain pairs were excluded from this statistic because they do not bear on the adversity-brain question. In the training set, the null distribution was generated by permuting the adversity view across training families (whole families moved together) and refitting the model. In the held-out set, the training weights were held fixed and the adversity rows were permuted, so held-out *p*-values describe the fixed training solution rather than a re-estimated one. Components 1-5 were tested with 10,000 permutations (minimum attainable *p* = .0001), as were the per-view canonical correlations. Each was scored on views deflated by the preceding components, with the held-out views deflated using the training loadings.

Stability selection^23^ refitted the sparse model (s = 0.4) in 10,000 subsamples, each drawing 70% of training families without replacement, and recorded the proportion of subsamples in which each feature received a non-zero component-1 weight. Following Lett et al.^21^, features selected in at least 90% of subsamples were considered stably selected. Structural loadings (the correlation of each feature with its own view’s variate) were estimated over 10,000 bootstrap resamples of training families. In each resample the non-sparse (s = 1) component-1 model was refitted, and its loadings were sign-aligned to the full training reference solution by dot product. The reported loading is the bootstrap mean with a 95% percentile interval. The two-sided bootstrap *p*-value was twice the smaller of the proportions of resamples above and below zero, with a floor of 1/10,000, and Benjamini-Hochberg false discovery rate (FDR) correction^24^ was applied within each view. Notably, the weight shows whether the sparse solution used a feature; selection frequency shows how reliably it did so across subsamples; and a loading shows how strongly the feature covaries with the view’s variate, whether or not the feature carries weight. Therefore, a feature can load significantly on the variate without being selected, because correlated features can stand in for it.

Canonical variates have arbitrary sign. Throughout, the adversity variate is oriented so that higher scores indicate greater adversity (all stably selected adversity features load positively). In the tables, each brain view’s loadings are reported in the orientation of that view’s own variate. The sign of its correlation with the adversity variate is stated in each table note.

#### Non-Sparse Companion Variates

The sparse model was used for feature selection, and a companion model refitted with no sparsity (s=1) on the same eight views and the same training/held-out split supplied all subject-level scores. Its weights were estimated in the training set and applied unchanged to held-out participants. The resulting adversity, SA, and rsFC scores were used for the held-out scatterplots (main-text Fig. 1d), the year-6 outcome models, the mediation analyses, the threat-versus-deprivation composites, and the within-family decomposition.

#### Supplementary Brain-Side Analyses

##### Two-view models and cumulative-risk comparison

To estimate each imaging modality’s association with adversity without competition from the other views, we fitted two-view sparse models (the adversity view against one imaging view, s = 0.4) in the training set and evaluated them in the held-out set. The same two-view estimator underlies the cumulative risk comparison, the global-versus-regional decomposition, the cross-modal convergence analysis, and the baseline-only, one-per-family, and held-out-site sensitivity analyses. For the cumulative risk comparison^25^, the learned weighting was compared with an equal-weight sum of the same 104 residualized and standardized features, which formed a coherent scale (Cronbach’s α = 0.82) and correlated *r* = 0.78 with the sparse adversity variate in held-out participants. Both predictors were related to the SA and rsFC views by two-view models fitted in the training set. The difference in held-out |*r*| was bootstrapped over 2,000 paired resamples of held-out families, with the same families serving both predictors in each draw.

##### Global versus regional surface area

Three analyses distinguished a global from a regional SA association. First, total SA and ICV were each regressed on the adversity variate (after both were residualized on age, sex, and site) in the training, held-out, and full samples, with family-clustered 95% CIs. Second, two-view models related adversity to four representations of SA: the 68 parcels as measured (the reference), total SA alone as a single feature, the 68 parcels with total SA residualized from each, and the 68 parcels with ICV residualized from each. The residualization coefficients were estimated in the training set. Retention was the held-out |*r*| of each representation as a percentage of the reference. Its 95% CI came from 2,000 bootstrap resamples of held-out families, with numerator and denominator recomputed on the same resample and the training weights held fixed. Third, regional stability selection was repeated in 2,000 subsamples of 70% of training families for the unadjusted and the total-SA-residualized parcels, using the same draws for both.

##### Cross-modal convergence

To test whether the SA and rsFC signatures reflect a common environmental gradient, a separate two-view model was fitted in the training set for each of the seven imaging views. The training weights were then used to score all participants, and the seven adversity-side scores were correlated with one another and with the joint adversity variate in held-out participants. The corresponding brain-side scores were correlated in the same way. Every adversity axis was oriented to the same deprivation anchor so that the signs of the correlations are interpretable. If two modalities pick out the same environmental gradient, their adversity-side scores should be nearly collinear, whatever the correlation between their brain-side scores.

##### Threat versus deprivation

Features were assigned *a priori* to threat and deprivation^26^. The primary tagging used only instruments whose assignment is uncontroversial. Threat comprised the KSADS traumatic-events items, caregiver- and youth-reported family conflict, neighborhood unsafety, and neighborhood crime (39 features). Deprivation comprised household income and education, financial hardship, neglect, low caregiver acceptance, low parental monitoring, area deprivation, child opportunity, social vulnerability, social mobility, and tree canopy (25 features). The remaining 40 features were left unassigned, including the Life Events Scale, whose items mix threat and loss and could not be allocated without item text; caregiver psychopathology; family substance history; and the environmental-hazard LED measures. A secondary tagging added the life events to threat and the hazards to deprivation (64 and 30 features). For each tagging, equal-weight composites (the mean of the residualized, standardized features) were correlated with the non-sparse SA and rsFC variates in held-out participants, both marginally and with the other composite partialled out. Separate two-view models were then fitted for threat-only and deprivation-only feature sets against SA and against rsFC, and the correlation of their held-out brain-side scores was compared with a null distribution of 100 random splits of the same pooled features into sets of the same two sizes.

##### Within-family decomposition

Within-family analyses used the 657 families with two or more members in the analytic sample (1,330 participants). For each non-sparse variate we estimated the sibling intraclass correlation (ICC) with a family-bootstrap 95% CI and a linear mixed-model cross-check. We then decomposed the association between the adversity variate and each brain variate into a within-family component (correlation of deviations from family means) and a between-family component (correlation of family means), with 1,000 family-bootstrap resamples.

#### Year-6 Outcome Models and Mediation

Each Year-6 outcome was regressed on a non-sparse variate by ordinary least squares with family-clustered robust standard errors. Covariates were age at the Year-6 visit, sex, site, and, for the CBCL^8^ and NIH Toolbox^27^ outcomes, the baseline value of the outcome. Predictors and outcomes were *z*-scored within each model. Cognition was analyzed as NIH Toolbox uncorrected standard scores. BH-FDR correction was applied across outcomes within each predictor and domain, where the psychopathology and cognition outcomes formed one family and the five cardiometabolic outcomes another. For the incremental test, both brain variates (SA and rsFC) were added to the model containing the adversity variate. We evaluated the change in R^2^ and a two-degree-of-freedom cluster-robust Wald test of the brain block (FDR-corrected across outcomes within each domain), and we report each brain variate’s partial coefficient. The SA and rsFC variates were weakly correlated, so collinearity was negligible (variance inflation factors ≤ 1.11).

For mediation, the a-path regressed the SA variate on the adversity variate, and the b-path regressed the outcome on the SA variate adjusting for the adversity variate. Both paths used the covariates of the outcome model. The indirect effect (a x b) and the proportion mediated (indirect / total effect) were bootstrapped over 5,000 family-clustered resamples with percentile 95% CIs, and indirect-effect *p*-values were FDR-corrected across the five outcomes tested (the four outcomes with significant adversity associations plus glycated hemoglobin). A proportion mediated was not interpreted when its interval was unstable. Because the adversity and SA variates were both measured at or near baseline, the mediation model partitions a statistical association and does not establish temporal precedence of adversity over brain structure or a causal pathway. Two further baseline-adjustment analyses were run. The CBCL models were refitted without the baseline covariate, so that they describe year-6 symptom levels rather than change. The waist-to-height ratio model, which in the primary specification has no baseline covariate, was refitted with baseline waist-to-height ratio added, both on the primary rows and on the identical complete-case rows used for the adjusted model (below).

#### Sensitivity Analyses

Each sensitivity analysis changed one feature of the primary specification and held everything else constant, including the training-only residualization, which was re-estimated within each analysis. Retention was defined as the held-out |*r*| under the sensitivity specification as a percentage of the corresponding primary held-out |*r*|, with a 95% CI from 5,000 bootstrap resamples of held-out families (training weights fixed, one resample serving numerator and denominator). Retention above 100% indicates a larger estimate in the sensitivity sample, not necessarily a stronger underlying association.

*Baseline-only battery.* This analysis addresses whether the dimension depends on the two instruments collected after the imaging visit. It removed the 25 Life Events and 8 neglect items (104 to 71 features). *One participant per family.* This analysis addresses whether the solution depends on related participants. It retained the first-listed member of each family within each split (training 3,712, held-out 1,591). *Held-out sites.* This analysis addresses generalization across sites and scanners. The model was trained on 15 sites and evaluated on 6 entirely held-out sites (training 4,373, held-out 1,603). Because the held-out sites contribute no training rows, their site effects cannot be estimated from training data, so each held-out feature was centered within its site after the training age, sex, and motion coefficients were applied. For this analysis, retention intervals were also computed by resampling sites. For all three sensitivity analyses, the structural and functional associations were evaluated with two-view models for total SA and rsFC, and the year-6 cognition coefficients were re-estimated with each specification’s own refitted adversity variate. These three analyses did not refit the five-component joint model or its permutation tests.

*Site residualization of the adversity view.* This analysis addresses whether residualizing site from the adversity view removed meaningful environmental variance or shaped the solution. Site was removed from the adversity view’s covariate set (age and sex were still residualized), and all seven imaging views were unchanged. The full joint pipeline was rerun: eight-view SGCCA at s = 0.4, 10,000 permutations for each of the five components, and 10,000-draw stability selection. *Penalty (s = 0.7).* This analysis addresses whether conclusions depend on the chosen sparsity. The model was refitted at the cross-validation optimum with the blocks unchanged, and the same permutation tests were run; stability selection was not repeated at s = 0.7. For both analyses, weights, structural loadings, and variate scores were compared with the primary solution. *Baseline waist-to-height ratio.* This analysis addresses whether the prospective waist-to-height association reflects adiposity already present at baseline. It is described under the outcome models above. The intracranial volume analysis is part of the global-versus-regional decomposition.

### Supplementary Results

#### Representativeness of the Analytic Sample

Relative to the 5,884 baseline participants without usable data for every imaging view, the analytic sample (n = 5,976) was slightly older, more often female, more often White, and more socioeconomically advantaged (Table S6). The differences were small to modest in magnitude (Cramér’s *V* = 0.07 for sex to 0.17 for caregiver education; Cohen’s *d* = 0.21 for age), but all were statistically significant. The largest differences were in the proportions of Black participants and of households with income below $50,000, both lower in the analytic sample. These differences follow documented associations between scan quality and sociodemographic factors in ABCD^28^ and imply a restricted range of environmental disadvantage relative to the full cohort.

#### Component Structure and Modality Specificity

Component 1 was the only component whose adversity-brain statistic exceeded its permutation null in both the training (observed 0.0128, null mean 0.0022) and held-out (observed 0.0125, null mean 0.0023) sets (both *p* < .001). Components 2-5 did not exceed their null in either the training (*p*s ≥ .125) or the held-out (*p*s ≥ .432) set. Table S5 gives component 1 by view. Only the SA and rsFC variates were associated with the adversity variate in both splits. The SST variate reached nominal held-out significance (|*r*| = 0.052, *p* = .049) without a training association (|*r*| = 0.005), which does not meet the replication criterion. Cortical thickness retained 15 stably selected parcels in the sparse joint solution even though its variate did not replicate, and in two-view models CT reached a held-out |*r*| of 0.187, although its joint-model association did not replicate once SA and rsFC were modeled alongside it. The absence of replicating associations for the other views should therefore not be read as an absence of association between adversity and those measures.

#### Environmental Feature Selection

Table S1 lists the non-zero weight, selection frequency, and bootstrap structural loading of every adversity feature. Of the 104 features, 39 received non-zero weights in the primary sparse solution and 19 were stably selected (≥90% of subsamples). All 19 stable features also had non-zero weights in the full-training solution. By domain, the stable features comprised 8 neighborhood and built environment features, 8 household socioeconomic features, 2 caregiver psychopathology scales, and 1 interpersonal feature (witnessed household violence). The highest selection frequency among features that were not stably selected was 0.863 (youth-reported conflict item 6).

Because the adversity variate is a weighted combination of broadly and positively intercorrelated features, 85 of the 104 bootstrap loadings were FDR-significant, including 47 features with zero weight in the primary solution. Significance of a loading therefore indicates that a feature covaries with the socioeconomic dimension, not necessarily that it contributes to it independently of correlated features. We accordingly interpret features by the magnitude of their loadings and their selection frequency. Conversely, features that were not selected, including family conflict, neglect, caregiver acceptance, parental monitoring, and most life events, are not shown to be unrelated to brain structure or function. Within this multivariate model, their association with the brain was not reliably separable from the neighborhood and household socioeconomic features that carried the dimension.

#### Global Versus Regional Cortical Surface Area

Greater adversity was associated with smaller total SA in every sample (training *β* = -0.206, held-out *β* = -0.190, full sample *β* = -0.202; all *p* < .001) and with smaller ICV, less strongly (full sample *β* = -0.159 [-0.180, -0.138]). In two-view models (Table S7), total SA alone reached a held-out |*r*| of 0.225, which is 96.1% of the 68-parcel reference (|*r*| = 0.234). The interval included 100%. Residualizing ICV from each parcel retained 69.0%, and residualizing total SA retained 46.7%. Regional variance was substantially shared with global size: on average 42.2% of a parcel’s variance was explained by total SA. Adding the size-residualized regional axis to total SA increased the held-out variance explained in the adversity axis by ΔR^2^ = 0.0088.

At the regional level (Table S8), 12 of 68 parcels were stably selected in the unadjusted two-view model and none after total-SA residualization (maximum 0.784). In the joint eight-view model, 12 parcels were stably selected, all in frontal, temporal, or pericentral cortex. Every parcel loaded in the same direction on the SA variate, with bootstrap intervals excluding zero for all 68. This is the pattern expected when a single global factor underlies the regional features, and it is why regional loadings were not interpreted as evidence of localization.

#### Resting-State Connectivity

Table S9 lists all 78 rsFC features; Figure S1a shows the same loadings as a network matrix. 10 features were stably selected at 90% and 13 at 75%. Nine of the ten stable features involved a sensorimotor, auditory, or salience/cingulo-opercular community. The exception was dorsal attention-visual. No rsFC loading survived FDR correction (smallest *q* = .405), and only 8 of 78 bootstrap intervals excluded zero. Across bootstrap refits of the non-sparse model, the sign of individual edge loadings was not stable, even though the same edges were repeatedly selected by the sparse model. The reliable property of the rsFC signature is which network pairs enter the solution, not the sign or magnitude of any single pair. Because ABCD regresses the global signal before computing network correlations^14^, the signature reflects the relative configuration of connectivity rather than a global shift in connectivity strength.

#### Cross-Modal Convergence

When each modality was modeled against adversity separately, the adversity axes derived from SA and from rsFC were nearly collinear in held-out participants (*r* = 0.986; Table S10). Each correlated highly with the joint adversity variate (*r* = 0.957 and 0.938). The CT-, FA-, and MID-derived adversity axes were similarly aligned (*r* = 0.90-0.98 with the SA and rsFC axes). The SST- and ENB-derived axes, which had no held-out association with the brain, were not (*r* = 0.21 and 0.43 with the SA axis). On the brain side, the SA and rsFC scores were essentially uncorrelated (*r* = -0.053). Surface area and resting-state connectivity therefore index the same environmental gradient through largely independent brain features.

#### Threat Versus Deprivation

Under the primary tagging, the deprivation composite was more strongly associated with the SA variate than the threat composite was (held-out *r* = -0.172 vs -0.106; Table S11), and this difference widened when each composite was adjusted for the other (partial *r* = -0.147 vs -0.056). For rsFC, deprivation again exceeded threat (partial *r* = 0.074 vs 0.007). In separate two-view models, the deprivation-only feature set reached held-out |*r*| = 0.236 with SA and 0.189 with rsFC, compared with 0.113 and 0.119 for threat. The brain scores derived from threat-only and deprivation-only models were highly correlated (|*r*| = 0.948 for SA and 0.752 for rsFC). These correlations fell below every random split of the same features (null medians 0.993 and 0.958). That comparison should not be read as evidence of dissociable threat and deprivation neurobiology. Random splits place deprivation items in both halves, so their brain scores are nearly identical by construction, and any split that separates content will fall below that ceiling. The results were qualitatively unchanged under the secondary tagging (Table S11). Together they indicate a single brain axis that is more strongly associated with deprivation than with threat, not a separable threat-related brain dimension. The primary tagging leaves 40 features, including all 25 life events, unassigned.

#### Within-Family Decomposition

The adversity variate was almost entirely shared within families (ICC = 0.917 [0.893, 0.937]); only 8.3% of its variance lay within families (Table S12). The SA variate was moderately familial (ICC = 0.604) and the rsFC variate weakly so (ICC = 0.213). Among siblings, the adversity-SA association was carried by between-family differences (*r* = -0.266 [-0.334, -0.197]), and the within-family estimate was small, with an interval that included zero (*r* = -0.070 [-0.150, 0.005]). The rsFC association showed the same pattern (between *r* = 0.177; within *r* = 0.012).

These estimates cannot be used to separate environmental from familial influences. With less than a tenth of the exposure’s variance available within families, and only a few hundred families contributing, within-family estimators have little information about this dimension. A within-family null is therefore uninformative. Equally, the between-family association cannot exclude family level confounding, including heritable parental characteristics. The analysis establishes the structure of the exposure (a family level socioeconomic gradient) rather than the causal source of its brain association.

#### Year-6 Outcomes: Baseline Adjustment, Brain Increments, and Mediation

Table S13 gives the full year-6 results for all ten outcomes. Without the baseline covariate, the adversity variate predicted higher year-6 internalizing, externalizing, and total problems (*β* = 0.12-0.16, all *p* < .001; Table S14). With baseline symptoms adjusted, these associations were near zero. The dimension is therefore related to symptom levels already present at ages 9-10 years rather than to symptom change over the following six years. The baseline-adjusted waist-to-height ratio result is reported in Table S14. On the 3,095 participants with baseline waist-to-height ratio, the primary specification gave *β* = 0.214, identical to the primary estimate on all 3,101 participants (*β* = 0.214). Adding baseline waist-to-height ratio reduced the coefficient to *β* = 0.112 [0.080, 0.143], *p* < .001, a 48% attenuation on identical rows. A significant prospective association with waist-to-height ratio remained beyond baseline adiposity, although about half of the unadjusted association was already evident at ages 9-10 years.

In the incremental models, the brain block added significant prediction for crystallized and fluid cognition, glycated hemoglobin, diastolic blood pressure, and CBCL total problems (Table S13). For crystallized and fluid cognition, both brain variates contributed partial coefficients, SA positive and rsFC negative. Mediation through the SA variate was significant for fluid and crystallized cognition and diastolic blood pressure, and absent for waist-to-height ratio. The glycated hemoglobin proportion mediated is not interpreted because its total effect was close to zero and its interval was very wide. These statistical indirect effects are consistent with a pathway through cortical SA.

#### Baseline-Only Battery, One Participant per Family, and Held-Out Sites

Table S15 summarizes the three standard sensitivity analyses. Restricting the battery to baseline measures retained 98.5% of the held-out total-SA association and 96.9% of the rsFC association. Retaining one participant per family retained 101.9% and 99.4%. Training on 15 sites and testing on 6 held-out sites retained 100.5% of the rsFC association and yielded a larger held-out total-SA estimate (132.0%). In that analysis the held-out total-SA |*r*| exceeded the training |*r*| (0.297 vs 0.205), and the site-resampled interval was wide (95% CI 99.5%-176.5%), which reflects between-site heterogeneity rather than a stronger association. In all specifications, the adversity variate predicted lower Year-6 fluid and crystallized cognition in held-out participants (all *p* < .001).

#### Site Residualization of the Adversity View

When site was not residualized from the adversity view, the joint model again yielded a single replicating component (held-out adversity-brain statistic 0.0096, *p* < .001; components 2-5 null; Table S16), and SA and rsFC were again the only replicating views (held-out |*r*| = 0.178 and 0.177). The adversity weights were nearly identical to the primary weights (*r* = 0.992), and the same 39 features were non-zero. The 10 largest loadings were the same five neighborhood and five household socioeconomic features as in the primary model. Stability selection identified 18 stable adversity features, all among the primary 19; noise exposure fell just below threshold. Restoring between-site variance, however, meant that site explained 17.3% of the training adversity variate. The held-out SA and rsFC associations were modestly attenuated, retaining 85.0% [75.1, 95.0] and 93.2% [82.9, 104.4] of the primary estimates. The number of stably selected rsFC features fell from 10 to 6, although rsFC selection frequencies correlated r = 0.994 with the primary frequencies, so edges near the threshold moved to either side of it. Between-site environmental variance therefore adds noise with respect to the brain rather than a different dimension.

#### Penalty Sensitivity (L1 = 0.7)

At the cross-validation-optimal penalty (s = 0.7), one component again replicated (held-out statistic 0.0126, *p* < .001; components 2-5 did not replicate; Table S16), with SA (held-out |*r*| = 0.224) and rsFC (|*r*| = 0.173) as the only views replicating in both splits. Component 2 reached nominal significance in held-out data when the fixed training weights were applied (*p* = .004), but was not significant in training (*p* = .358) or when the model was refit (held-out *p* = .144), and therefore did not meet the prespecified replication criterion. Retention was 107.2% [99.3, 116.3] for SA and 90.7% [71.8, 112.5] for rsFC. The weaker penalty produced a far less sparse solution: 103 of 104 adversity features and 54 of 78 rsFC features had non-zero weights, compared with 39 and 21 at s = 0.4. The environmental profile was preserved. The adversity structural loadings correlated *r* = 0.979 with the primary loadings, the adversity variate scores correlated *r* = 0.958 in held-out participants, and the 10 largest loadings were again five neighborhood and five household socioeconomic features. The SA pattern was likewise preserved (loadings *r* = 0.978). The rsFC composition was less stable. Weights correlated only *r* = 0.40 and held-out variate scores *r* = 0.67 with the primary solution, and default-mode pairs (Auditory - Default mode, Cingulo-parietal - Default mode) entered the ten largest loadings. The s = 0.7 solution is not equally sparse and does not reproduce the primary rsFC composition edge for edge. It preserves the primary dimensionality, the modality specificity (SA and rsFC), and the socioeconomic and neighborhood interpretation of the environmental variate. The Year-6 outcome models, which use the non-sparse companion scores, do not depend on the penalty.

### Supplementary Tables

***Table S1.*** *The 104-Feature Adversity and Environmental Battery: Source, Coding, Selection, and Loadings*

| Feature | ABCD variable(s) | Coding | Missing (%) | Non-zero weight | Selection frequency | Loading [95% CI] | *q* |
| --- | --- | --- | --- | --- | --- | --- | --- |
| *Neighborhood and built environment (13 features)* | | | | | | | |
| *Neighborhood Safety and Crime Scale: Caregiver / youth; Baseline* | | | | | | | |
| Neighborhood unsafe (caregiver; mean of 3 items) | fc_p_nsc__ns_001-003 | Reversed; higher = less safe | 0.0 | Yes | **1.00** | 0.51 [0.46, 0.55] | .001 |
| Neighborhood unsafe (youth; 1 item) | fc_y_nsc__ns_003 | Reversed; higher = less safe | 0.1 | Yes | **1.00** | 0.36 [0.32, 0.40] | .002 |
| *Linked external data: Residential address; Baseline address* | | | | | | | |
| Area Deprivation Index (national percentile) | le_l_adi__addr1__national_prcnt | Higher = more deprived | 6.5 | Yes | **1.00** | 0.69 [0.64, 0.74] | .002 |
| Child Opportunity Index (national score) | le_l_coi__addr1__coi__total__national_score | Reversed; higher = lower opportunity | 8.4 | Yes | **1.00** | 0.73 [0.68, 0.78] | .002 |
| Social Vulnerability Index (percentile) | le_l_svi__addr1__total_prcntile | Higher = more vulnerable | 8.4 | Yes | **1.00** | 0.68 [0.62, 0.73] | .002 |
| Social mobility | le_l_socmob__addr1__kfrpp_mean | Reversed; higher = lower mobility | 8.4 | Yes | **1.00** | 0.70 [0.65, 0.75] | .001 |
| Nitrogen dioxide, annual mean (2016) | le_l_no2__addr1__no2_mean__2016 | Higher = more exposure | 4.8 | — | 0.11 | 0.18 [0.12, 0.23] | .002 |
| Fine particulate matter (PM2.5), annual mean (2016) | le_l_pm25__addr1__pm25_mean__2016 | Higher = more exposure | 4.8 | — | 0.44 | 0.16 [0.11, 0.21] | .002 |
| Lead exposure risk (percentile) | le_l_leadrisk__addr1__homerisk_prcnt | Higher = more risk | 4.8 | — | 0.14 | 0.29 [0.23, 0.34] | .001 |
| Noise, 24-h mean sound level | le_l_noise__addr1__24hmean__total_soundlvl | Higher = more noise | 4.7 | Yes | **0.98** | 0.32 [0.26, 0.39] | .001 |
| Tree canopy (% cover) | le_l_nlcd__addr1__treecn_prcnt | Reversed; higher = less canopy | 4.7 | Yes | **1.00** | 0.33 [0.28, 0.39] | .001 |
| Alcohol-outlet density (2016) | le_l_densalc__addr1__alc_density__2016 | Higher = more outlets | 7.2 | — | 0.01 | 0.06 [0.00, 0.12] | .049 |
| Crime (summed reported counts) | le_l_crime__* counts (summed) | Higher = more crime | 4.8 | — | 0.00 | 0.08 [0.04, 0.12] | .002 |
| *Household socioeconomic (9 features)* | | | | | | | |
| *Demographics questionnaire: Caregiver; Baseline* | | | | | | | |
| Partner education | ab_p_demo__edu__prtnr_001 | Reversed; higher = less education | 16.3 | Yes | **1.00** | 0.41 [0.37, 0.46] | .001 |
| Caregiver education | ab_p_demo__edu__slf_001 | Reversed; higher = less education | 0.1 | Yes | **1.00** | 0.59 [0.55, 0.63] | .002 |
| Household income | ab_p_demo__income__hhold_001 | Reversed; higher = lower income | 7.1 | Yes | **1.00** | 0.65 [0.62, 0.68] | .001 |
| *Demographics: financial hardship: Caregiver; Baseline* | | | | | | | |
| Could not afford food | ab_p_demo__exp__fam_001 | 1 = hardship | 0.4 | Yes | **1.00** | 0.46 [0.40, 0.52] | .001 |
| Without telephone service (unaffordable) | ab_p_demo__exp__fam_002 | 1 = hardship | 0.2 | Yes | **1.00** | 0.45 [0.39, 0.50] | .002 |
| Did not pay full rent or mortgage | ab_p_demo__exp__fam_003 | 1 = hardship | 0.3 | Yes | **1.00** | 0.47 [0.42, 0.53] | .002 |
| Evicted from home | ab_p_demo__exp__fam_004 | 1 = hardship | 0.2 | Yes | 0.54 | 0.22 [0.16, 0.28] | .002 |
| Utilities shut off for non-payment | ab_p_demo__exp__fam_005 | 1 = hardship | 0.2 | Yes | **1.00** | 0.33 [0.28, 0.39] | .002 |
| Did not seek needed medical care (cost) | ab_p_demo__exp__fam_006 | 1 = hardship | 0.2 | Yes | **0.99** | 0.30 [0.24, 0.36] | .002 |
| *Caregiver psychopathology and substance use (10 features)* | | | | | | | |
| *Adult Self-Report, DSM-oriented scales: Caregiver; Baseline* | | | | | | | |
| ADHD problems | mh_p_asr__dsm__adhd_sum | Raw sum; higher = more | 0.0 | — | 0.33 | 0.30 [0.21, 0.37] | .002 |
| Antisocial personality problems | mh_p_asr__dsm__antsoc_sum | Raw sum; higher = more | 0.0 | Yes | **1.00** | 0.36 [0.28, 0.43] | .002 |
| Anxiety problems | mh_p_asr__dsm__anx_sum | Raw sum; higher = more | 0.0 | Yes | 0.57 | 0.31 [0.24, 0.38] | .002 |
| Avoidant personality problems | mh_p_asr__dsm__avoid_sum | Raw sum; higher = more | 0.0 | — | 0.02 | 0.26 [0.18, 0.33] | .002 |
| Depressive problems | mh_p_asr__dsm__dep_sum | Raw sum; higher = more | 0.0 | — | 0.29 | 0.33 [0.24, 0.41] | .002 |
| Somatic problems | mh_p_asr__dsm__somat_sum | Raw sum; higher = more | 0.0 | Yes | **1.00** | 0.35 [0.29, 0.41] | .002 |
| *Family History Assessment: Caregiver; Baseline* | | | | | | | |
| Father, alcohol problem | mh_p_famhx__alc__fath_001___1-7 | 1 = any problem reported | 0.0 | — | 0.19 | 0.19 [0.13, 0.25] | .001 |
| Father, drug problem | mh_p_famhx__drg__fath_001___1-7 | 1 = any problem reported | 0.0 | — | 0.34 | 0.25 [0.19, 0.31] | .001 |
| Mother, alcohol problem | mh_p_famhx__alc__moth_001___1-7 | 1 = any problem reported | 0.0 | Yes | 0.46 | 0.16 [0.11, 0.22] | .001 |
| Mother, drug problem | mh_p_famhx__drg__moth_001___1-7 | 1 = any problem reported | 0.0 | Yes | 0.75 | 0.19 [0.12, 0.25] | .001 |
| *Interpersonal and life events (72 features)* | | | | | | | |
| *KSADS traumatic-events screener: Caregiver; Baseline* | | | | | | | |
| Traumatic event 1 | mh_p_ksads__ptsd__screen_001___1 | 1 = event endorsed | 2.8 | — | 0.27 | 0.11 [0.06, 0.16] | .002 |
| Traumatic event 2 | mh_p_ksads__ptsd__screen_001___2 | 1 = event endorsed | 2.8 | — | 0.00 | 0.01 [-0.04, 0.06] | .711 |
| Traumatic event 3 | mh_p_ksads__ptsd__screen_001___3 | 1 = event endorsed | 2.8 | — | 0.02 | 0.08 [0.03, 0.14] | .006 |
| Traumatic event 4 | mh_p_ksads__ptsd__screen_001___4 | 1 = event endorsed | 2.8 | — | 0.00 | 0.01 [-0.03, 0.06] | .636 |
| Traumatic event 5 | mh_p_ksads__ptsd__screen_001___5 | 1 = event endorsed | 2.8 | — | 0.00 | 0.05 [-0.01, 0.14] | .203 |
| Traumatic event 6 | mh_p_ksads__ptsd__screen_001___6 | 1 = event endorsed | 2.8 | — | 0.02 | 0.07 [0.01, 0.15] | .020 |
| Traumatic event 7 | mh_p_ksads__ptsd__screen_001___7 | 1 = event endorsed | 2.8 | — | 0.13 | 0.14 [0.08, 0.22] | .002 |
| Traumatic event 8 | mh_p_ksads__ptsd__screen_001___8 | 1 = event endorsed | 2.8 | — | 0.30 | 0.11 [0.01, 0.23] | .038 |
| Traumatic event 9 | mh_p_ksads__ptsd__screen_001___9 | 1 = event endorsed | 2.8 | — | 0.00 | 0.11 [0.02, 0.24] | .020 |
| Traumatic event 10 | mh_p_ksads__ptsd__screen_001___10 | 1 = event endorsed | 2.8 | — | 0.01 | 0.12 [0.03, 0.22] | .010 |
| Traumatic event 11 | mh_p_ksads__ptsd__screen_001___11 | 1 = event endorsed | 2.8 | — | 0.00 | 0.03 [-0.03, 0.10] | .513 |
| Traumatic event 12 | mh_p_ksads__ptsd__screen_001___12 | 1 = event endorsed | 2.8 | — | 0.00 | 0.08 [0.02, 0.16] | .010 |
| Traumatic event 13: witnessed household violence | mh_p_ksads__ptsd__screen_001___13 | 1 = event endorsed | 2.8 | Yes | **0.97** | 0.28 [0.22, 0.34] | .001 |
| Traumatic event 14 | mh_p_ksads__ptsd__screen_001___14 | 1 = event endorsed | 2.8 | — | 0.00 | 0.08 [0.00, 0.17] | .049 |
| Traumatic event 15 | mh_p_ksads__ptsd__screen_001___15 | 1 = event endorsed | 2.8 | — | 0.00 | 0.07 [0.01, 0.16] | .042 |
| Traumatic event 16 | mh_p_ksads__ptsd__screen_001___16 | 1 = event endorsed | 2.8 | — | 0.01 | 0.04 [-0.02, 0.11] | .197 |
| Traumatic event 17 | mh_p_ksads__ptsd__screen_001___17 | 1 = event endorsed | 2.6 | — | 0.17 | 0.18 [0.14, 0.23] | .002 |
| Traumatic event 18 | mh_p_ksads__ptsd__screen_001___18 | 1 = event endorsed | 2.1 | — | 0.00 | -0.01 [-0.05, 0.04] | .765 |
| *Family Environment Scale, conflict: Caregiver; Baseline* | | | | | | | |
| Conflict item 1 | fc_p_fes__confl_001 | 1 = conflict endorsed | 0.0 | — | 0.00 | 0.11 [0.04, 0.17] | .004 |
| Conflict item 2 | fc_p_fes__confl_002 | 1 = conflict endorsed | 0.0 | — | 0.04 | 0.12 [0.06, 0.18] | .002 |
| Conflict item 3 | fc_p_fes__confl_003 | 1 = conflict endorsed | 0.0 | — | 0.31 | 0.19 [0.13, 0.25] | .002 |
| Conflict item 4 | fc_p_fes__confl_004 | 1 = conflict endorsed | 0.0 | Yes | 0.52 | 0.16 [0.10, 0.22] | .002 |
| Conflict item 5 | fc_p_fes__confl_005 | 1 = conflict endorsed | 0.0 | Yes | 0.54 | 0.20 [0.13, 0.26] | .002 |
| Conflict item 6 | fc_p_fes__confl_006 | 1 = conflict endorsed | 0.0 | Yes | 0.68 | 0.20 [0.14, 0.26] | .002 |
| Conflict item 7 | fc_p_fes__confl_007 | 1 = conflict endorsed | 0.0 | — | 0.00 | 0.05 [0.00, 0.10] | .039 |
| Conflict item 8 | fc_p_fes__confl_008 | 1 = conflict endorsed | 0.0 | — | 0.02 | 0.19 [0.13, 0.25] | .002 |
| Conflict item 9 | fc_p_fes__confl_009 | 1 = conflict endorsed | 0.1 | Yes | 0.63 | -0.07 [-0.12, -0.01] | .026 |
| *Family Environment Scale, conflict: Youth; Baseline* | | | | | | | |
| Conflict item 1 | fc_y_fes__confl_001 | 1 = conflict endorsed | 0.1 | Yes | 0.71 | 0.18 [0.12, 0.24] | .002 |
| Conflict item 2 | fc_y_fes__confl_002 | 1 = conflict endorsed | 0.1 | Yes | 0.53 | 0.12 [0.07, 0.17] | .001 |
| Conflict item 3 | fc_y_fes__confl_003 | 1 = conflict endorsed | 0.1 | Yes | 0.60 | 0.15 [0.09, 0.20] | .002 |
| Conflict item 4 | fc_y_fes__confl_004 | 1 = conflict endorsed | 0.1 | — | 0.00 | 0.09 [0.04, 0.15] | .003 |
| Conflict item 5 | fc_y_fes__confl_005 | 1 = conflict endorsed | 0.1 | Yes | 0.52 | 0.15 [0.10, 0.21] | .002 |
| Conflict item 6 | fc_y_fes__confl_006 | 1 = conflict endorsed | 0.1 | Yes | 0.86 | 0.16 [0.10, 0.21] | .002 |
| Conflict item 7 | fc_y_fes__confl_007 | 1 = conflict endorsed | 0.1 | Yes | 0.51 | 0.12 [0.07, 0.17] | .002 |
| Conflict item 8 | fc_y_fes__confl_008 | 1 = conflict endorsed | 0.1 | — | 0.35 | 0.12 [0.07, 0.17] | .002 |
| Conflict item 9 | fc_y_fes__confl_009 | 1 = conflict endorsed | 0.1 | — | 0.05 | 0.09 [0.04, 0.13] | .002 |
| *Life Events Scale: Caregiver; Year 1* | | | | | | | |
| Life event 1 | mh_p_ple_001 | 1 = event occurred | 4.3 | — | 0.08 | 0.12 [0.07, 0.16] | .001 |
| Life event 2 | mh_p_ple_002 | 1 = event occurred | 4.3 | — | 0.00 | 0.05 [0.00, 0.10] | .045 |
| Life event 3 | mh_p_ple_003 | 1 = event occurred | 4.3 | — | 0.00 | 0.08 [0.03, 0.13] | .002 |
| Life event 4 | mh_p_ple_004 | 1 = event occurred | 4.3 | — | 0.00 | 0.00 [-0.05, 0.05] | .964 |
| Life event 5 | mh_p_ple_005 | 1 = event occurred | 4.3 | — | 0.00 | -0.01 [-0.06, 0.04] | .655 |
| Life event 6 | mh_p_ple_006 | 1 = event occurred | 4.3 | — | 0.00 | -0.02 [-0.07, 0.02] | .280 |
| Life event 7 | mh_p_ple_007 | 1 = event occurred | 4.3 | — | 0.19 | 0.22 [0.16, 0.28] | .001 |
| Life event 8 | mh_p_ple_008 | 1 = event occurred | 4.3 | — | 0.00 | 0.13 [0.07, 0.20] | .002 |
| Life event 9 | mh_p_ple_009 | 1 = event occurred | 4.3 | — | 0.00 | 0.00 [-0.04, 0.05] | .901 |
| Life event 10 | mh_p_ple_010 | 1 = event occurred | 4.3 | — | 0.00 | -0.01 [-0.06, 0.03] | .604 |
| Life event 11 | mh_p_ple_011 | 1 = event occurred | 4.3 | — | 0.00 | 0.09 [0.03, 0.14] | .005 |
| Life event 12 | mh_p_ple_012 | 1 = event occurred | 4.3 | — | 0.01 | 0.09 [0.03, 0.15] | .006 |
| Life event 13 | mh_p_ple_013 | 1 = event occurred | 4.3 | — | 0.00 | 0.09 [0.03, 0.15] | .007 |
| Life event 14 | mh_p_ple_014 | 1 = event occurred | 4.3 | Yes | 0.80 | 0.25 [0.18, 0.32] | .001 |
| Life event 15 | mh_p_ple_015 | 1 = event occurred | 4.3 | — | 0.00 | 0.11 [0.05, 0.17] | .002 |
| Life event 16 | mh_p_ple_016 | 1 = event occurred | 4.3 | — | 0.00 | 0.09 [0.04, 0.14] | .002 |
| Life event 17 | mh_p_ple_017 | 1 = event occurred | 4.3 | — | 0.03 | 0.11 [0.06, 0.17] | .002 |
| Life event 18 | mh_p_ple_018 | 1 = event occurred | 4.3 | — | 0.01 | 0.12 [0.05, 0.18] | .002 |
| Life event 19 | mh_p_ple_019 | 1 = event occurred | 4.3 | Yes | 0.56 | 0.21 [0.13, 0.28] | .002 |
| Life event 20 | mh_p_ple_020 | 1 = event occurred | 4.3 | — | 0.00 | 0.01 [-0.04, 0.07] | .711 |
| Life event 21 | mh_p_ple_021 | 1 = event occurred | 4.3 | — | 0.01 | -0.02 [-0.08, 0.04] | .621 |
| Life event 22 | mh_p_ple_022 | 1 = event occurred | 4.3 | Yes | 0.67 | 0.24 [0.16, 0.31] | .001 |
| Life event 23 | mh_p_ple_023 | 1 = event occurred | 4.3 | — | 0.04 | 0.06 [0.01, 0.12] | .038 |
| Life event 24 | mh_p_ple_024 | 1 = event occurred | 4.3 | Yes | 0.53 | -0.04 [-0.10, 0.01] | .181 |
| Life event 25 | mh_p_ple_025 | 1 = event occurred | 4.3 | — | 0.21 | -0.01 [-0.06, 0.04] | .737 |
| *Multidimensional Neglectful Behavior Scale: Youth; Year 3* | | | | | | | |
| Educational support item 1 | fc_y_mnbs__edusupp_001 | Reversed; higher = more neglect | 9.5 | — | 0.00 | 0.02 [-0.03, 0.07] | .402 |
| Supervision item 1 | fc_y_mnbs__superv_001 | Reversed; higher = more neglect | 9.5 | Yes | 0.43 | 0.09 [0.04, 0.14] | .002 |
| Educational support item 2 | fc_y_mnbs__edusupp_002 | Reversed; higher = more neglect | 9.4 | — | 0.04 | 0.03 [-0.01, 0.08] | .141 |
| Supervision item 2 | fc_y_mnbs__superv_002 | Reversed; higher = more neglect | 9.6 | — | 0.00 | 0.05 [-0.00, 0.10] | .077 |
| Educational support item 3 | fc_y_mnbs__edusupp_003 | Reversed; higher = more neglect | 9.5 | — | 0.00 | 0.08 [0.03, 0.13] | .004 |
| Supervision item 3 | fc_y_mnbs__superv_003 | Reversed; higher = more neglect | 9.5 | — | 0.25 | 0.13 [0.08, 0.18] | .002 |
| Supervision item 4 | fc_y_mnbs__superv_004 | Reversed; higher = more neglect | 9.7 | — | 0.03 | 0.07 [0.02, 0.13] | .010 |
| Supervision item 5 | fc_y_mnbs__superv_005 | Reversed; higher = more neglect | 9.5 | — | 0.01 | -0.03 [-0.08, 0.02] | .290 |
| *CRPBI, acceptance: Youth; Baseline* | | | | | | | |
| Caregiver 1 acceptance (mean of 5 items) | fc_y_crpbi__cg1_002-006 | Reversed; higher = less warmth | 0.1 | — | 0.00 | 0.11 [0.05, 0.16] | .002 |
| Caregiver 2 acceptance (mean of 5 items) | fc_y_crpbi__cg2_002-006 | Reversed; higher = less warmth | 6.6 | — | 0.00 | 0.09 [0.04, 0.14] | .002 |
| *Parental Monitoring Scale: Youth; Baseline* | | | | | | | |
| Parental monitoring (mean of 5 items) | fc_y_pm_001-005 | Reversed; higher = less monitoring | 0.1 | Yes | 0.71 | 0.17 [0.12, 0.23] | .002 |

**Note.** Features are grouped by descriptive domain and instrument (italic rows give the instrument, informant, and assessment). Baseline = ABCD baseline visit (ages 9-10 years); Year 1 and Year 3 = first administration at the year-1 and year-3 follow-ups. Every feature is coded so that higher values indicate greater adversity; ‘Reversed’ marks protective constructs whose scores were reversed. Missing = percentage of the analytic sample (N = 5,976) imputed with the training-set median. Non-zero weight = non-zero component-1 weight in the primary sparse model (L1 = 0.4) fitted to all training participants (39 features). Selection frequency = proportion of 10,000 subsamples of 70% of training families in which the feature received a non-zero weight; bold marks stable selection (≥ 0.90; 19 features). Loading = structural loading (correlation of the feature with the adversity variate), bootstrap mean and 95% percentile interval over 10,000 resamples of training families; the adversity variate is oriented so that positive loadings indicate greater adversity. *q* = Benjamini-Hochberg FDR-corrected bootstrap *p* within the view (85 of 104 < .05). Significant loadings do not imply selection, and non-selection does not imply the absence of an association. Item wording for the Life Events, KSADS, Family Environment, and neglect items is given in the ABCD data dictionary under the listed variable names. ADHD = attention-deficit/hyperactivity disorder; CRPBI = Children’s Report of Parental Behavior Inventory; FDR = false discovery rate; KSADS = Kiddie Schedule for Affective Disorders and Schizophrenia; LED = linked external data; PM2.5 = particulate matter ≤ 2.5 µm.

***Table S2.*** *Harmonized MRI Acquisition Parameters by Scanner Platform*

| Parameter | Siemens | Philips | GE |
| --- | --- | --- | --- |
| *Platform* | | | |
| Scanner model (software) | Prisma / Prisma Fit (VE11B-C) | Achieva dStream / Ingenia | MR750 (DV25-26) |
| *3D T1w* | | | |
| Matrix | 256 × 256 | 256 × 256 | 256 × 256 |
| Slices | 176 | 225 | 208 |
| FOV (mm) | 256 × 256 | 256 × 240 | 256 × 256 |
| % FOV phase | 100% | 93.75% | 100% |
| Resolution (mm) | 1.0 × 1.0 × 1.0 | 1.0 × 1.0 × 1.0 | 1.0 × 1.0 × 1.0 |
| TR (ms) | 2500 | 6.31 | 2500 |
| TE (ms) | 2.88 | 2.9 | 2 |
| TI (ms) | 1060 | 1060 | 1060 |
| Flip angle (deg) | 8 | 8 | 8 |
| Parallel imaging | 2x | 1.5 × 2.2 | 2x |
| Multiband | Off | Off | Off |
| Acquisition time | 07:12 | 05:38 | 06:09 |
| *3D T2w* | | | |
| Matrix | 256 × 256 | 256 × 256 | 256 × 256 |
| Slices | 176 | 256 | 208 |
| FOV (mm) | 256 × 256 | 256 × 256 | 256 × 256 |
| Resolution (mm) | 1.0 × 1.0 × 1.0 | 1.0 × 1.0 × 1.0 | 1.0 × 1.0 × 1.0 |
| TR (ms) | 3200 | 2500 | 3200 |
| TE (ms) | 565 | 251.6 | 60 |
| TI (ms) | N/A | N/A | N/A |
| Flip angle (deg) | Variable | 90 | Variable |
| Parallel imaging | 2x | 1.5 × 2.0 | 2x |
| Multiband | Off | Off | Off |
| Acquisition time | 06:35 | 02:53 | 05:50 |
| *dMRI* | | | |
| Matrix | 140 × 140 | 140 × 140 | 140 × 140 |
| Slices | 81 | 81 | 81 |
| FOV (mm) | 240 × 240 | 240 × 240 | 240 × 240 |
| Resolution (mm) | 1.7 × 1.7 × 1.7 | 1.7 × 1.7 × 1.7 | 1.7 × 1.7 × 1.7 |
| TR (ms) | 4100 | 5300 | 4100 |
| TE (ms) | 88 | 89 | 81.9 |
| Flip angle (deg) | 90 | 78 | 77 |
| Parallel imaging | Off | Off | Off |
| Multiband factor | 3 | 3 | 3 |
| Partial Fourier / half-scan factor | 6/8 | 0.6 | 5.5/8 |
| Diffusion directions (total) | 96 | 96 | 96 |
| Directions per shell (s/mm²) | 6 @ b=500; 15 @ b=1000; 15 @ b=2000; 60 @ b=3000 | 6 @ b=500; 15 @ b=1000; 15 @ b=2000; 60 @ b=3000 | 6 @ b=500; 15 @ b=1000; 15 @ b=2000; 60 @ b=3000 |
| Acquisition time | 07:31 | 09:14 | 07:30 |
| *fMRI (rest and task)* | | | |
| Matrix | 90 × 90 | 90 × 90 | 90 × 90 |
| Slices | 60 | 60 | 60 |
| FOV (mm) | 216 × 216 | 216 × 216 | 216 × 216 |
| Resolution (mm) | 2.4 × 2.4 × 2.4 | 2.4 × 2.4 × 2.4 | 2.4 × 2.4 × 2.4 |
| TR (ms) | 800 | 800 | 800 |
| TE (ms) | 30 | 30 | 30 |
| Flip angle (deg) | 52 | 52 | 52 |
| Parallel imaging | Off | Off | Off |
| Multiband factor | 6 | 6 | 6 |
| Partial Fourier / half-scan factor | Off | 0.9 | Off |

**Note.** Values are from Casey et al.^13^ (Table 2), which match the ABCD protocol sheet; scanner models follow Hagler et al.^14^. The same fMRI parameters were used for resting-state and task runs (per-run time not tabulated in the source): rest, up to four 5-min runs; MID, two runs of 50 trials (5:42 each); SST, two runs of 180 trials (349 s each); ENB, two runs of eight blocks (160 trials). dMRI = diffusion MRI; ENB = emotional n-back; FOV = field of view; MID = monetary incentive delay; SST = stop-signal task; T1w/T2w = T1-/T2-weighted; TE = echo time; TI = inversion time; TR = repetition time.

***Table S3.*** *Neuroimaging Feature Sets and Native-Unit Descriptive Statistics*

| View | Features | Definition | Motion covariate | *n* passing QC | Native-unit summary, *M* (*SD*) |
| --- | --- | --- | --- | --- | --- |
| Cortical thickness (CT) | 68 | Desikan-Killiany parcels (34 per hemisphere), mm | None | 11,554 | 2.771 (0.074) mm (mean parcel) |
| Cortical surface area (SA) | 68 | Desikan-Killiany parcels, mm^2^ | None | 11,554 | 190,289 (17,661) mm² (total) |
| Fractional anisotropy (FA) | 37 | AtlasTrack: 17 tracts per hemisphere, corpus callosum, forceps major and minor | dMRI head motion | 10,404 | 0.461 (0.027) (mean FA); motion 1.264 (0.329) mm |
| Resting-state connectivity (rsFC) | 78 | 12 within- and 66 between-network mean correlations, 12 Gordon communities | Mean FD (rest) | 9,554 | FD 0.076 (0.019) mm (censored) |
| MID task | 84 | Large reward vs neutral anticipation; 68 cortical + 16 subcortical/cerebellar | Mean FD (MID) | 9,256 | FD 0.242 (0.227) mm |
| Stop-signal task (SST) | 84 | Correct stop vs correct go; same parcels | Mean FD (SST) | 8,099 | FD 0.250 (0.256) mm |
| Emotional n-back (ENB) | 84 | Emotional vs neutral faces; same parcels | Mean FD (ENB) | 7,819 | FD 0.272 (0.279) mm |

**Note.** All features are ABCD Release 7.0 tabulated DAIRC derivatives. *n* passing QC = baseline participants with usable data for that view before intersection; the analytic sample (N = 5,976) is the complete-case intersection of all views and the adversity view. Native-unit summaries are computed in the analytic sample before residualization; intracranial volume, used only in the global-versus-regional analyses, was 1,502,716 (139,546) mm^3^. Resting-state FD is computed over retained frames after censoring and task FD over all frames, so the two are not comparable; the dMRI value is a separate head-motion estimate. The motion covariate was residualized from that view only. dMRI = diffusion MRI; FD = framewise displacement; MID = monetary incentive delay; QC = quality control.

***Table S4.*** *Penalty Selection by Five-Fold Family-Grouped Cross-Validation*

| L1 penalty (*s*) | Mean criterion | *SD* across folds | *SE* | Optimum minus *s*, mean (*SE*) |
| --- | --- | --- | --- | --- |
| 0.1 | 0.0000 | 0.0000 | 0.0000 | 0.0838 (0.0061) |
| 0.2 | 0.0658 | 0.0083 | 0.0037 | 0.0180 (0.0070) |
| 0.3 | 0.0780 | 0.0130 | 0.0058 | 0.0058 (0.0079) |
| 0.4 (primary) | 0.0822 | 0.0112 | 0.0050 | 0.0016 (0.0056) |
| 0.5 | 0.0814 | 0.0110 | 0.0049 | 0.0024 (0.0020) |
| 0.7 (optimum) | 0.0838 | 0.0137 | 0.0061 | — |

**Note.** Criterion = mean absolute correlation between the adversity variate and each of the seven brain variates in the held-out fold, averaged over five folds of the training set grouped by family (siblings never straddle folds). The last column is the paired across-fold difference between the optimum (*s* = 0.7) and each value. At *s* = 0.1 the constraint admits no non-zero solution, so the criterion is zero. The primary value *s* = 0.4 was fixed for sparsity and interpretability because the criterion is flat between 0.4 and 0.7.

***Table S5.*** *Component 1 by View: Sparsity, Stability, and Training and Held-Out Associations*

| View | Features | Non-zero weights | Stable (≥ 0.90) | Training \|*r*\| | *p* | Held-out \|*r*\| | *p* | Two-view held-out \|*r*\| | Non-sparse held-out \|*r*\| |
| --- | --- | --- | --- | --- | --- | --- | --- | --- | --- |
| Adversity | 104 | 39 | 19 | — | — | — | — | — | — |
| Surface area | 68 | 22 | 12 | 0.210 | <.001 | 0.209 | <.001 | 0.234 | 0.226 |
| Resting-state connectivity | 78 | 21 | 10 | 0.195 | <.001 | 0.190 | <.001 | 0.207 | 0.138 |
| Cortical thickness | 68 | 19 | 15 | 0.041 | .375 | 0.000 | .993 | 0.187 | 0.023 |
| Fractional anisotropy | 37 | 7 | 0 | 0.050 | .407 | 0.030 | .293 | 0.067 | 0.052 |
| MID task | 84 | 27 | 0 | 0.044 | .373 | 0.034 | .094 | 0.063 | 0.034 |
| Stop-signal task | 84 | 25 | 0 | 0.005 | .908 | 0.052 | .049 | 0.004 | 0.055 |
| Emotional n-back | 84 | 30 | 0 | 0.031 | .550 | 0.055 | .079 | 0.034 | 0.034 |

**Note.** Joint eight-view SGCCA, component 1, L1 = 0.4. |*r*| = absolute correlation between the adversity variate and the view’s variate. Training and held-out *p*-values are from 10,000 permutations (training: adversity view permuted across families and model refitted; held-out: training weights fixed, adversity rows permuted); the minimum attainable *p* is .0001. Stable = features selected in ≥ 90% of 10,000 subsamples. Two-view = adversity view modeled against that imaging view alone (L1 = 0.4). Non-sparse = companion model at L1 = 1 used for subject-level analyses; its held-out |*r*| values are the main-text subject-level estimates. Training *N* = 4,186; held-out *N* = 1,790. MID = monetary incentive delay; SGCCA = sparse generalized canonical correlation analysis.

***Table S6.*** *Analytic Sample Versus Excluded Baseline Participants*

| Characteristic | Included  (*n* = 5,976) | Excluded  (*n* = 5,884) | Test, effect size | *p* |
| --- | --- | --- | --- | --- |
| *Age at baseline (years)* | | | | |
| M (SD) | 10.02 (0.63) | 9.89 (0.62) | *t* = 11.57, *d* = 0.21 | <.001 |
| *Sex* | | | | |
| Female | 3,062 (51.2) | 2,613 (44.4) | χ²(1) = 55.1, *V* = 0.07 | <.001 |
| Male | 2,914 (48.8) | 3,271 (55.6) |  |  |
| *Race/ethnicity* | | | | |
| White | 3,525 (59.0) | 2,660 (45.2) | χ^2^ (4) = 335.0, *V* = 0.17 | <.001 |
| Hispanic | 1,130 (18.9) | 1,313 (22.3) |  |  |
| Black | 606 (10.1) | 1,209 (20.5) |  |  |
| Asian | 121 (2.0) | 109 (1.9) |  |  |
| Other | 593 (9.9) | 593 (10.1) |  |  |
| *Household income* | | | | |
| < $50,000 | 1,266 (22.8) | 1,955 (37.0) | χ^2^ (2) = 290.1, *V* = 0.16 | <.001 |
| $50,000-$99,999 | 1,609 (29.0) | 1,458 (27.6) |  |  |
| $100,000 and greater | 2,679 (48.2) | 1,876 (35.5) |  |  |
| *Highest caregiver education* | | | | |
| Up to high school (no diploma) | 203 (3.4) | 390 (6.6) | χ^2^ (4) = 345.0, *V* = 0.17 | <.001 |
| High school diploma/GED | 404 (6.8) | 727 (12.4) |  |  |
| Some college | 1,349 (22.6) | 1,725 (29.4) |  |  |
| Bachelor's degree | 1,647 (27.6) | 1,362 (23.2) |  |  |
| Graduate or professional degree | 2,370 (39.7) | 1,669 (28.4) |  |  |

**Note.** Excluded = the remaining ABCD Release 7.0 baseline participants (cohort *N* = 11,860) without usable data for all seven imaging views. Values are *n* (%) unless stated; percentages exclude missing values (household income missing for 422 included and 595 excluded participants; caregiver education for 3 and 11; race/ethnicity for 1 included participant). Race/ethnicity is the ABCD five-level cohort variable, in which Hispanic ethnicity takes precedence over racial categories. Tests are Welch’s *t* test for age and Pearson χ^2^ tests on the full level distribution for categorical variables. *d* = Cohen’s *d*; GED = General Educational Development credential; *V* = Cramér’s *V*.

***Table S7.*** *Global Versus Regional Decomposition of the Surface-Area Association*

| SA representation | Training \|*r*\| / β | Held-out \|*r*\| / β | Retention, % [95% CI] | Non-zero adversity weights | Stable parcels (≥ 0.90) |
| --- | --- | --- | --- | --- | --- |
| *Two-view models: adversity view vs surface-area representation* | | | | | |
| 68 parcels as measured (reference) | 0.252 | 0.234 | 100 (reference) | 41 | 12 |
| Total SA alone (1 feature) | 0.231 | 0.225 | 96.1 [89.5, 102.5] | 36 | — |
| 68 parcels, ICV residualized | 0.172 | 0.161 | 69.0 [52.9, 85.2] | 29 | Not run |
| 68 parcels, total SA residualized | 0.138 | 0.109 | 46.7 [27.8, 66.1] | 39 | 0 |
| *Adversity variate predicting global size: standardized β [95% CI]* | | | | | |
| Total SA | -0.206 [-0.232, -0.180] | -0.190 [-0.230, -0.150] | Full sample: -0.202 [-0.224, -0.180] | — | — |
| Intracranial volume | -0.162 [-0.187, -0.136] | -0.153 [-0.191, -0.114] | Full sample: -0.159 [-0.180, -0.138] | — | — |

**Note.** Upper panel: two-view sparse models (L1 = 0.4) fitted in training families (*N* = 4,186) and evaluated in held-out families (*N* = 1,790). Retention = held-out |*r*| as a percentage of the 68-parcel reference, with a 95% percentile interval from 2,000 bootstrap resamples of held-out families (numerator and denominator recomputed on the same resample; training weights fixed). Residualization coefficients were estimated in training data. Stable parcels = parcels selected in ≥ 90% of 2,000 subsamples of 70% of training families (same draws for both rows); stability selection was not run for the ICV-residualized parcels. Lower panel: adversity variate predicting total SA and ICV, both residualized on age, sex, and site; standardized β with family-clustered 95% CI (training *N* = 4,186, held-out *N* = 1,790, full *N* = 5,976; all *p* < .001). Total SA correlated *r* = 0.767 with ICV. ICV = intracranial volume; SA = surface area.

***Table S8.*** *Regional Surface Area Loadings and Selection Before and After Total Surface Area Adjustment*

| Region | Hemi | Loading [95% CI] | Joint-model selection | Two-view selection, unadjusted | Two-view selection, total SA residualized |
| --- | --- | --- | --- | --- | --- |
| Middle temporal | RH | 0.76 [0.75, 0.78] | **1.00** | **1.00** | 0.76 |
| Inferior temporal | RH | 0.70 [0.68, 0.71] | **0.92** | **1.00** | 0.78 |
| Fusiform | LH | 0.69 [0.67, 0.71] | 0.00 | **1.00** | 0.42 |
| Inferior temporal | LH | 0.67 [0.65, 0.69] | 0.34 | **1.00** | 0.75 |
| Precentral | RH | 0.67 [0.65, 0.69] | **1.00** | **1.00** | 0.47 |
| Precentral | LH | 0.70 [0.68, 0.72] | **1.00** | **1.00** | 0.11 |
| Fusiform | RH | 0.70 [0.68, 0.71] | 0.00 | **0.99** | 0.14 |
| Precuneus | LH | 0.70 [0.68, 0.72] | 0.24 | **0.96** | 0.03 |
| Precuneus | RH | 0.71 [0.69, 0.73] | 0.75 | **0.95** | 0.01 |
| Middle temporal | LH | 0.70 [0.68, 0.72] | 0.89 | **0.92** | 0.21 |
| Superior frontal | RH | 0.77 [0.75, 0.78] | **1.00** | **0.91** | 0.01 |
| Superior parietal | RH | 0.61 [0.59, 0.63] | 0.07 | **0.91** | 0.09 |
| Superior parietal | LH | 0.63 [0.61, 0.65] | 0.13 | 0.85 | 0.09 |
| Postcentral | LH | 0.72 [0.70, 0.74] | **1.00** | 0.84 | 0.01 |
| Caudal middle frontal | LH | 0.60 [0.58, 0.63] | 0.87 | 0.79 | 0.35 |
| Lateral orbitofrontal | LH | 0.76 [0.74, 0.77] | 0.72 | 0.75 | 0.02 |
| Caudal middle frontal | RH | 0.60 [0.57, 0.62] | **0.99** | 0.58 | 0.26 |
| Pars orbitalis | RH | 0.61 [0.59, 0.63] | 0.01 | 0.44 | 0.13 |
| Frontal pole | LH | 0.59 [0.57, 0.62] | 0.62 | 0.40 | 0.20 |
| Lateral occipital | LH | 0.58 [0.56, 0.60] | 0.00 | 0.39 | 0.34 |
| Lateral occipital | RH | 0.58 [0.55, 0.60] | 0.00 | 0.35 | 0.36 |
| Medial orbitofrontal | RH | 0.73 [0.72, 0.75] | 0.03 | 0.33 | 0.05 |
| Pars orbitalis | LH | 0.65 [0.63, 0.67] | 0.71 | 0.31 | 0.13 |
| Superior frontal | LH | 0.77 [0.76, 0.79] | **1.00** | 0.22 | 0.49 |
| Frontal pole | RH | 0.57 [0.55, 0.60] | 0.01 | 0.20 | 0.17 |
| Inferior parietal | RH | 0.59 [0.57, 0.61] | 0.01 | 0.08 | 0.11 |
| Postcentral | RH | 0.68 [0.66, 0.69] | 0.65 | 0.05 | 0.15 |
| Rostral middle frontal | LH | 0.70 [0.68, 0.72] | **1.00** | 0.04 | 0.08 |
| Rostral ant. cingulate | LH | 0.67 [0.65, 0.69] | 0.78 | 0.04 | 0.03 |
| Lateral orbitofrontal | RH | 0.72 [0.70, 0.73] | 0.05 | 0.04 | 0.12 |
| Rostral middle frontal | RH | 0.68 [0.66, 0.70] | **1.00** | 0.02 | 0.06 |
| Superior temporal | RH | 0.76 [0.75, 0.78] | **1.00** | 0.01 | 0.72 |
| Inferior parietal | LH | 0.59 [0.56, 0.61] | 0.01 | 0.00 | 0.10 |
| Rostral ant. cingulate | RH | 0.58 [0.56, 0.60] | 0.00 | 0.00 | 0.00 |
| Pars opercularis | RH | 0.52 [0.49, 0.54] | 0.00 | 0.00 | 0.07 |
| Superior temporal | LH | 0.73 [0.71, 0.74] | **1.00** | 0.00 | 0.64 |
| Medial orbitofrontal | LH | 0.68 [0.66, 0.70] | 0.00 | 0.00 | 0.23 |
| Insula | LH | 0.66 [0.64, 0.68] | 0.69 | 0.00 | 0.66 |
| Insula | RH | 0.64 [0.62, 0.66] | 0.42 | 0.00 | 0.15 |
| Posterior cingulate | RH | 0.63 [0.60, 0.65] | 0.00 | 0.00 | 0.67 |
| Supramarginal | RH | 0.60 [0.57, 0.62] | 0.00 | 0.00 | 0.24 |
| Posterior cingulate | LH | 0.60 [0.58, 0.62] | 0.00 | 0.00 | 0.09 |
| Isthmus cingulate | LH | 0.58 [0.56, 0.60] | 0.00 | 0.00 | 0.10 |
| Paracentral | RH | 0.57 [0.55, 0.59] | 0.59 | 0.00 | 0.10 |
| Paracentral | LH | 0.57 [0.54, 0.59] | 0.14 | 0.00 | 0.08 |
| Transverse temporal | RH | 0.57 [0.54, 0.59] | 0.10 | 0.00 | 0.69 |
| Supramarginal | LH | 0.57 [0.54, 0.59] | 0.01 | 0.00 | 0.05 |
| Isthmus cingulate | RH | 0.55 [0.52, 0.57] | 0.00 | 0.00 | 0.47 |
| Temporal pole | LH | 0.53 [0.51, 0.56] | 0.02 | 0.00 | 0.04 |
| Temporal pole | RH | 0.52 [0.50, 0.55] | 0.00 | 0.00 | 0.36 |
| Banks sup. temporal sulcus | RH | 0.52 [0.50, 0.55] | 0.00 | 0.00 | 0.07 |
| Pars triangularis | LH | 0.51 [0.49, 0.54] | 0.00 | 0.00 | 0.39 |
| Pars opercularis | LH | 0.51 [0.49, 0.54] | 0.00 | 0.00 | 0.03 |
| Lingual | RH | 0.51 [0.48, 0.54] | 0.00 | 0.00 | 0.37 |
| Lingual | LH | 0.51 [0.48, 0.53] | 0.00 | 0.00 | 0.20 |
| Caudal ant. cingulate | LH | 0.50 [0.48, 0.53] | 0.00 | 0.00 | 0.00 |
| Transverse temporal | LH | 0.50 [0.47, 0.53] | 0.01 | 0.00 | 0.65 |
| Cuneus | RH | 0.49 [0.46, 0.52] | 0.00 | 0.00 | 0.28 |
| Pars triangularis | RH | 0.48 [0.45, 0.50] | 0.00 | 0.00 | 0.76 |
| Caudal ant. cingulate | RH | 0.48 [0.45, 0.50] | 0.00 | 0.00 | 0.03 |
| Parahippocampal | LH | 0.48 [0.45, 0.50] | 0.00 | 0.00 | 0.04 |
| Banks sup. temporal sulcus | LH | 0.47 [0.44, 0.50] | 0.00 | 0.00 | 0.15 |
| Parahippocampal | RH | 0.47 [0.44, 0.49] | 0.00 | 0.00 | 0.01 |
| Cuneus | LH | 0.43 [0.40, 0.46] | 0.00 | 0.00 | 0.23 |
| Pericalcarine | RH | 0.38 [0.35, 0.41] | 0.00 | 0.00 | 0.44 |
| Pericalcarine | LH | 0.35 [0.32, 0.39] | 0.00 | 0.00 | 0.21 |
| Entorhinal | RH | 0.31 [0.27, 0.34] | 0.00 | 0.00 | 0.71 |
| Entorhinal | LH | 0.31 [0.27, 0.34] | 0.00 | 0.00 | 0.59 |

**Note.** Loading = structural loading on the SA variate of the joint model (bootstrap mean and 95% percentile interval, 10,000 resamples of training families; all 68 FDR *q* < .001). All loadings are positive: the SA variate indexes larger area throughout, and it correlates negatively with the adversity variate (non-sparse held-out *r* = -0.226), so greater adversity corresponds to smaller area in every parcel. Joint-model selection = proportion of 10,000 subsamples of the eight-view model with a non-zero weight. Two-view selection = proportion of 2,000 subsamples of the adversity-versus-SA model, before and after total SA was residualized from each parcel. Bold marks ≥ 0.90. Rows are ordered by unadjusted two-view selection, then loading. Hemi = hemisphere (LH/RH); SA = surface area; FDR = false discovery rate.

***Table S9.*** *Resting-State Network Pair Loadings and Stability Selection (78 Features)*

| Network pair | Type | Loading [95% CI] | *q* | Selection frequency |
| --- | --- | --- | --- | --- |
| Sensorimotor hand - Sensorimotor mouth | Between | 0.32 [-0.15, 0.58] | .461 | **0.979** |
| Auditory - Cingulo-opercular | Between | -0.03 [-0.53, 0.46] | .929 | **0.978** |
| Auditory - Salience | Between | -0.31 [-0.63, 0.25] | .579 | **0.971** |
| Sensorimotor hand | Within | 0.35 [-0.03, 0.57] | .405 | **0.969** |
| Frontoparietal - Sensorimotor hand | Between | -0.35 [-0.62, 0.16] | .458 | **0.962** |
| Dorsal attention - Visual | Between | -0.20 [-0.40, 0.09] | .461 | **0.957** |
| Salience - Sensorimotor hand | Between | -0.22 [-0.59, 0.33] | .630 | **0.951** |
| Auditory - Frontoparietal | Between | -0.24 [-0.57, 0.27] | .582 | **0.929** |
| Cingulo-parietal - Sensorimotor hand | Between | -0.32 [-0.49, -0.02] | .405 | **0.910** |
| Frontoparietal - Sensorimotor mouth | Between | -0.27 [-0.52, 0.13] | .461 | **0.901** |
| Cingulo-opercular - Sensorimotor hand | Between | -0.04 [-0.42, 0.36] | .896 | 0.894 |
| Salience - Sensorimotor mouth | Between | -0.18 [-0.49, 0.26] | .630 | 0.779 |
| Cingulo-opercular | Within | 0.10 [-0.48, 0.54] | .839 | 0.768 |
| Auditory - Sensorimotor hand | Between | 0.37 [0.03, 0.58] | .405 | 0.660 |
| Cingulo-parietal - Salience | Between | 0.17 [-0.18, 0.39] | .582 | 0.614 |
| Auditory - Visual | Between | 0.23 [-0.30, 0.59] | .630 | 0.568 |
| Visual | Within | -0.13 [-0.50, 0.35] | .814 | 0.484 |
| Retrosplenial-temporal - Visual | Between | -0.20 [-0.47, 0.20] | .582 | 0.483 |
| Default mode - Sensorimotor mouth | Between | -0.31 [-0.49, -0.01] | .405 | 0.454 |
| Auditory - Sensorimotor mouth | Between | 0.24 [-0.02, 0.44] | .405 | 0.450 |
| Cingulo-parietal - Dorsal attention | Between | -0.25 [-0.41, 0.14] | .405 | 0.357 |
| Auditory - Default mode | Between | -0.40 [-0.58, -0.05] | .405 | 0.317 |
| Retrosplenial-temporal - Salience | Between | -0.04 [-0.42, 0.43] | .896 | 0.290 |
| Dorsal attention - Ventral attention | Between | 0.04 [-0.28, 0.33] | .880 | 0.274 |
| Retrosplenial-temporal | Within | -0.04 [-0.29, 0.24] | .880 | 0.248 |
| Dorsal attention - Retrosplenial-temporal | Between | -0.20 [-0.36, 0.04] | .405 | 0.245 |
| Salience - Ventral attention | Between | -0.24 [-0.45, 0.09] | .451 | 0.227 |
| Cingulo-parietal - Frontoparietal | Between | 0.25 [-0.02, 0.40] | .405 | 0.180 |
| Cingulo-opercular - Retrosplenial-temporal | Between | -0.13 [-0.58, 0.54] | .821 | 0.175 |
| Ventral attention - Visual | Between | 0.17 [-0.27, 0.48] | .664 | 0.121 |
| Cingulo-parietal - Sensorimotor mouth | Between | -0.31 [-0.46, -0.03] | .405 | 0.105 |
| Default mode - Ventral attention | Between | 0.03 [-0.32, 0.36] | .896 | 0.101 |
| Dorsal attention - Sensorimotor hand | Between | -0.10 [-0.31, 0.17] | .670 | 0.083 |
| Cingulo-opercular - Frontoparietal | Between | 0.01 [-0.15, 0.21] | .947 | 0.077 |
| Default mode - Dorsal attention | Between | -0.06 [-0.42, 0.36] | .880 | 0.073 |
| Cingulo-opercular - Sensorimotor mouth | Between | 0.08 [-0.30, 0.36] | .823 | 0.071 |
| Auditory - Retrosplenial-temporal | Between | -0.10 [-0.54, 0.49] | .846 | 0.064 |
| Dorsal attention | Within | 0.06 [-0.24, 0.33] | .839 | 0.060 |
| Default mode - Retrosplenial-temporal | Between | 0.33 [-0.11, 0.50] | .405 | 0.043 |
| Cingulo-opercular - Salience | Between | 0.13 [-0.33, 0.42] | .630 | 0.034 |
| Default mode - Salience | Between | -0.14 [-0.41, 0.18] | .579 | 0.032 |
| Cingulo-parietal - Default mode | Between | 0.39 [-0.10, 0.56] | .405 | 0.029 |
| Dorsal attention - Salience | Between | 0.29 [0.00, 0.47] | .405 | 0.028 |
| Cingulo-opercular - Cingulo-parietal | Between | -0.25 [-0.46, 0.25] | .458 | 0.024 |
| Auditory - Ventral attention | Between | -0.32 [-0.50, -0.02] | .405 | 0.023 |
| Cingulo-opercular - Dorsal attention | Between | 0.28 [-0.16, 0.47] | .405 | 0.021 |
| Ventral attention | Within | -0.02 [-0.27, 0.25] | .896 | 0.018 |
| Cingulo-opercular - Ventral attention | Between | -0.26 [-0.44, 0.02] | .405 | 0.018 |
| Cingulo-opercular - Default mode | Between | -0.24 [-0.51, 0.23] | .471 | 0.017 |
| Auditory - Dorsal attention | Between | 0.21 [-0.18, 0.40] | .451 | 0.013 |
| Frontoparietal - Salience | Between | 0.18 [-0.03, 0.33] | .405 | 0.013 |
| Cingulo-parietal - Retrosplenial-temporal | Between | 0.07 [-0.22, 0.29] | .735 | 0.009 |
| Sensorimotor hand - Visual | Between | 0.05 [-0.35, 0.42] | .889 | 0.009 |
| Retrosplenial-temporal - Sensorimotor mouth | Between | -0.14 [-0.36, 0.24] | .582 | 0.008 |
| Default mode - Sensorimotor hand | Between | -0.27 [-0.46, 0.04] | .405 | 0.007 |
| Dorsal attention - Frontoparietal | Between | 0.05 [-0.15, 0.22] | .817 | 0.007 |
| Sensorimotor mouth | Within | 0.06 [-0.09, 0.19] | .630 | 0.004 |
| Auditory - Cingulo-parietal | Between | -0.25 [-0.42, 0.00] | .405 | 0.004 |
| Sensorimotor hand - Ventral attention | Between | -0.02 [-0.17, 0.16] | .880 | 0.004 |
| Cingulo-parietal | Within | 0.13 [-0.04, 0.24] | .409 | 0.003 |
| Sensorimotor mouth - Visual | Between | 0.09 [-0.10, 0.23] | .582 | 0.002 |
| Retrosplenial-temporal - Ventral attention | Between | 0.16 [-0.16, 0.38] | .582 | 0.001 |
| Frontoparietal - Retrosplenial-temporal | Between | 0.12 [-0.02, 0.25] | .405 | 0.001 |
| Cingulo-parietal - Visual | Between | -0.21 [-0.35, -0.00] | .405 | 0.001 |
| Auditory | Within | 0.06 [-0.17, 0.27] | .814 | 0.001 |
| Cingulo-opercular - Visual | Between | 0.10 [-0.29, 0.41] | .814 | 0.001 |
| Cingulo-parietal - Ventral attention | Between | 0.11 [-0.09, 0.27] | .461 | 0.000 |
| Frontoparietal | Within | 0.11 [-0.13, 0.28] | .582 | 0.000 |
| Frontoparietal - Ventral attention | Between | -0.01 [-0.22, 0.22] | .941 | 0.000 |
| Default mode | Within | 0.17 [-0.23, 0.47] | .604 | 0.000 |
| Salience - Visual | Between | 0.13 [-0.14, 0.34] | .582 | 0.000 |
| Default mode - Visual | Between | 0.09 [-0.16, 0.31] | .670 | 0.000 |
| Salience | Within | 0.09 [-0.07, 0.22] | .579 | 0.000 |
| Dorsal attention - Sensorimotor mouth | Between | 0.03 [-0.13, 0.19] | .870 | 0.000 |
| Default mode - Frontoparietal | Between | 0.03 [-0.20, 0.22] | .872 | 0.000 |
| Frontoparietal - Visual | Between | 0.01 [-0.17, 0.19] | .898 | 0.000 |
| Retrosplenial-temporal - Sensorimotor hand | Between | -0.04 [-0.27, 0.22] | .870 | 0.000 |
| Sensorimotor mouth - Ventral attention | Between | -0.14 [-0.28, 0.04] | .405 | 0.000 |

**Note.** Features are mean Fisher-*z* correlations within or between the 12 Gordon communities, computed by ABCD after global-signal regression. Selection frequency = proportion of 10,000 subsamples of 70% of training families with a non-zero weight; bold marks stable selection (≥ 0.90; 10 features; 13 at ≥ 0.75). Loading = structural loading on the rsFC variate (bootstrap mean and 95% percentile interval, 10,000 resamples of training families, non-sparse refits sign-aligned to the reference); the rsFC variate correlates positively with the adversity variate (non-sparse held-out *r* = 0.138), so positive loadings indicate higher connectivity with greater adversity. Because individual edge loadings changed sign across bootstrap refits, no loading survived FDR correction (*q* = Benjamini-Hochberg-corrected bootstrap *p*), and 8 of 78 intervals exclude zero; the reliable property is which pairs are selected. Rows are ordered by selection frequency. rsFC = resting-state functional connectivity; FDR = false discovery rate.

***Table S10.*** *Cross-Modal Convergence of Single Modality Adversity and Brain Axes*

| Axis | SA | rsFC | CT | FA | MID | SST | ENB | Joint |
| --- | --- | --- | --- | --- | --- | --- | --- | --- |
| *A. Adversity-side scores* | | | | | | | | |
| SA | — |  |  |  |  |  |  |  |
| rsFC | 0.99 | — |  |  |  |  |  |  |
| CT | 0.98 | 0.98 | — |  |  |  |  |  |
| FA | 0.97 | 0.96 | 0.96 | — |  |  |  |  |
| MID | 0.90 | 0.92 | 0.90 | 0.88 | — |  |  |  |
| SST | 0.21 | 0.17 | 0.21 | 0.24 | 0.12 | — |  |  |
| ENB | 0.43 | 0.42 | 0.39 | 0.42 | 0.29 | 0.02 | — |  |
| Joint (8-view variate) | 0.96 | 0.94 | 0.92 | 0.92 | 0.83 | 0.22 | 0.54 | — |
| *B. Brain-side scores* | | | | | | | | |
| SA | — |  |  |  |  |  |  |  |
| rsFC | -0.05 | — |  |  |  |  |  |  |
| CT | 0.02 | -0.10 | — |  |  |  |  |  |
| FA | -0.04 | 0.03 | -0.11 | — |  |  |  |  |
| MID | -0.02 | 0.00 | 0.00 | -0.02 | — |  |  |  |
| SST | -0.03 | 0.06 | -0.07 | -0.01 | -0.05 | — |  |  |
| ENB | -0.03 | -0.03 | 0.01 | 0.02 | -0.00 | -0.01 | — |  |
| Joint (8-view variate) | -0.94 | 0.05 | -0.03 | 0.05 | 0.03 | 0.02 | 0.02 | — |

**Note.** Held-out correlations (*N* = 1,790) among the scores of seven two-view sparse models (adversity view vs each imaging view, L1 = 0.4, weights fitted in training families) and the joint eight-view variate. Panel A correlates the adversity-side scores, oriented to a common deprivation anchor; panel B correlates the corresponding brain-side scores. Joint = the non-sparse joint-model adversity variate (panel A) or SA variate (panel B). Held-out two-view canonical |*r*|: SA 0.234; rsFC 0.207; CT 0.187; FA 0.067; MID 0.063; SST 0.004; ENB 0.034. The joint SA brain variate correlates -0.94 with the two-view SA brain score because of an arbitrary sign difference. CT = cortical thickness; ENB = emotional n-back; FA = fractional anisotropy; MID = monetary incentive delay; rsFC = resting-state functional connectivity; SA = surface area; SST = stop-signal task.

***Table S11.*** *Threat Versus Deprivation Composites and Their Brain Associations*

| Brain measure | Threat *r* | Threat partial *r* | Deprivation *r* | Deprivation partial *r* | Threat-only two-view \|*r*\| | Deprivation-only two-view \|*r*\| | Brain-score \|*r*\| | Random-split null median [5th, 95th] |
| --- | --- | --- | --- | --- | --- | --- | --- | --- |
| *Primary tagging (threat 39 features, deprivation 25 features)* | | | | | | | | |
| Surface area | -0.106 | -0.056 | -0.172 | -0.147 | 0.113 | 0.236 | 0.948 | 0.993 [0.982, 0.997] |
| Resting-state connectivity | 0.031 | 0.007 | 0.080 | 0.074 | 0.119 | 0.189 | 0.752 | 0.958 [0.879, 0.987] |
| *Secondary tagging (threat 64 features, deprivation 30 features)* | | | | | | | | |
| Surface area | -0.097 | -0.058 | -0.155 | -0.134 | 0.122 | 0.235 | 0.966 | 0.993 [0.977, 0.997] |
| Resting-state connectivity | 0.039 | 0.016 | 0.086 | 0.079 | 0.126 | 0.189 | 0.903 | 0.961 [0.842, 0.987] |

**Note.** Held-out participants (*N* = 1,790). Composite columns: correlations of the equal-weight threat and deprivation composites with the non-sparse SA and rsFC variates, marginal and with the other composite partialled out (the SA variate indexes larger area, so negative values indicate smaller area with greater exposure). Two-view columns: held-out canonical |*r*| of separate sparse models (L1 = 0.4) using only the threat or only the deprivation features. Brain-score |*r*| = correlation between the brain-side scores of those two models; the null distribution is 100 random splits of the same pooled features into sets of the same two sizes. Random splits place deprivation items in both halves, forcing brain-score correlations near 1, so an observed value below the null reflects the content-separating split rather than dissociable neurobiology. Primary tagging: threat = KSADS traumatic events, family conflict (caregiver and youth), neighborhood unsafety, crime; deprivation = income, education, financial hardship, neglect, low acceptance, low monitoring, area deprivation, child opportunity, social vulnerability, social mobility, tree canopy; unassigned = life events, caregiver psychopathology, family substance history, environmental hazards. The secondary tagging adds life events to threat and hazards to deprivation. rsFC = resting-state functional connectivity; SA = surface area.

***Table S12.*** *Within-Family Decomposition in Sibling Families*

| Variate | ICC(1)  [95% CI] | Within-family variance (%) | *r* with adversity, pooled [95% CI] | Between-family *r*  [95% CI] | Within-family *r*  [95% CI] |
| --- | --- | --- | --- | --- | --- |
| Adversity variate | 0.917 [0.893, 0.937] | 8.3 | — | — | — |
| Surface-area variate | 0.604 [0.540, 0.658] | 39.6 | -0.238 [-0.294, -0.177] | -0.266 [-0.334, -0.197] | -0.070 [-0.150, 0.005] |
| Resting-state variate | 0.213 [0.137, 0.282] | 78.7 | 0.134 [0.074, 0.188] | 0.177 [0.101, 0.248] | 0.012 [-0.068, 0.085] |

**Note.** Sibling subsample: 1,330 participants in 657 families with two or more members in the analytic sample. Non-sparse variate scores. ICC(1) = intraclass correlation with family-bootstrap 95% CI (a linear mixed-model estimate agreed to the third decimal: 0.918 for adversity). Between-family *r* = correlation of family means; within-family *r* = correlation of deviations from family means; CIs from 1,000 family-bootstrap resamples. Because the adversity variate has little within-family variance, within-family estimates are uninformative about its association with the brain; they are not evidence against it. Signs follow the variate orientation (the SA variate indexes larger area). ICC = intraclass correlation; SA = surface area.

***Table S13.*** *Year-6 Outcomes: Adversity Associations, Brain Increments, and Surface Area Mediation*

| Outcome | *n* | Adversity β [95% CI] | *q* | Brain block Δ*R*² | *q* | SA partial β | rsFC partial β | Indirect effect via SA [95% CI] | *q* | % mediated [95% CI] |
| --- | --- | --- | --- | --- | --- | --- | --- | --- | --- | --- |
| *Cognition and psychopathology (baseline value of the outcome covaried)* | | | | | | | | | | |
| Fluid cognition | 3,718 | -0.173 [-0.201, -0.144] | <.001 | 0.0044 | <.001 | **0.034** | **-0.053** | -0.0090 [-0.0149, -0.0030] | .005 | 5.2 [1.7, 8.9] |
| Crystallized cognition | 3,851 | -0.095 [-0.118, -0.072] | <.001 | 0.0079 | <.001 | **0.058** | **-0.063** | -0.0113 [-0.0158, -0.0071] | .002 | 11.8 [7.2, 18.1] |
| CBCL internalizing | 4,846 | 0.017 [-0.010, 0.044] | .268 | 0.0007 | .109 | -0.005 | **-0.028** | Not tested |  |  |
| CBCL externalizing | 4,849 | 0.025 [-0.004, 0.054] | .151 | 0.0007 | .109 | -0.019 | -0.023 | Not tested |  |  |
| CBCL total problems | 4,846 | -0.001 [-0.029, 0.027] | .932 | 0.0013 | .025 | -0.007 | **-0.037** | Not tested |  |  |
| *Cardiometabolic health (no baseline covariate)* | | | | | | | | | | |
| Waist-to-height ratio | 3,101 | 0.214 [0.176, 0.252] | <.001 | 0.0003 | .787 | -0.008 | 0.014 | 0.0022 [-0.0055, 0.0096] | .587 | — |
| Systolic blood pressure | 3,061 | 0.025 [-0.010, 0.060] | .204 | 0.0010 | .311 | -0.018 | 0.025 | Not tested |  |  |
| Diastolic blood pressure | 3,061 | 0.115 [0.079, 0.151] | <.001 | 0.0028 | .024 | **-0.054** | -0.001 | 0.0120 [0.0040, 0.0204] | .005 | 10.4 [3.4, 20.1] |
| Total cholesterol | 1,565 | -0.026 [-0.071, 0.020] | .275 | 0.0003 | .787 | 0.018 | 0.002 | Not tested |  |  |
| Glycated hemoglobin | 2,068 | 0.058 [0.001, 0.115] | .080 | 0.0047 | .024 | **-0.061** | 0.029 | 0.0133 [0.0049, 0.0226] | .004 | Not interpretable |

**Note.** Year-6 outcomes regressed on non-sparse variates by OLS with family-clustered robust SEs, covarying age at the Year-6 visit, sex, and site (and the baseline outcome for cognition and CBCL); predictors and outcomes z-scored within model. Cognition = NIH Toolbox uncorrected standard scores. Adversity β is from the model with the adversity variate as the only predictor of interest. Brain block = both brain variates added to that model; Δ*R*² and cluster-robust Wald test (2 *df*); partial β in bold where FDR *q* < .05 (the SA variate indexes larger area). Mediation: indirect effect of the adversity variate through the SA variate (5,000 family-clustered bootstraps), tested for the five outcomes shown; % mediated reported only where the indirect effect was significant; the glycated hemoglobin proportion is not interpretable because the total effect was near zero and its interval extended to 98%. *q* = Benjamini-Hochberg FDR within predictor and domain (cognition and psychopathology; cardiometabolic) or across the five mediation tests. Blood pressure was modeled as the *z* scores supplied in the scored ABCD blood-pressure file. BP = blood pressure; CBCL = Child Behavior Checklist; FDR = false discovery rate; NIHTB = NIH Toolbox; OLS = ordinary least squares; rsFC = resting-state functional connectivity; SA = surface area.

***Table S14.*** *Baseline Adjustment of Year-6 Psychopathology and Waist-to-Height Ratio Associations*

| Outcome / model | Baseline covaried | *n* | Adversity β [95% CI] | *p* | *q* | *R*² |
| --- | --- | --- | --- | --- | --- | --- |
| *A. Year-6 CBCL, with and without the baseline score covaried* | | | | | | |
| Internalizing | Yes | 4,846 | 0.017 [-0.010, 0.044] | .215 | .268 | 0.236 |
| Internalizing | No | 4,847 | 0.117 [0.085, 0.148] | <.001 | <.001 | 0.050 |
| Externalizing | Yes | 4,849 | 0.025 [-0.004, 0.054] | .091 | .151 | 0.263 |
| Externalizing | No | 4,850 | 0.155 [0.121, 0.189] | <.001 | <.001 | 0.033 |
| Total problems | Yes | 4,846 | -0.001 [-0.029, 0.027] | .932 | .932 | 0.308 |
| Total problems | No | 4,847 | 0.155 [0.122, 0.189] | <.001 | <.001 | 0.036 |
| *B. Year-6 waist-to-height ratio, with and without baseline waist-to-height ratio covaried* | | | | | | |
| Primary model, all rows | No | 3,101 | 0.214 [0.176, 0.252] | <.001 | — | 0.106 |
| Primary model, complete-case rows | No | 3,095 | 0.214 [0.176, 0.252] | <.001 | — | 0.106 |
| Baseline-adjusted model | Yes | 3,095 | 0.112 [0.080, 0.143] | <.001 | — | 0.410 |

**Note.** Standardized coefficients for the non-sparse adversity variate; OLS with family-clustered robust SEs, covarying age at the Year-6 visit, sex, and site. Panel A: baseline-adjusted rows are the primary models; *q* = Benjamini-Hochberg FDR within each specification’s outcome family (the five cognition and psychopathology outcomes for the baseline-adjusted models; the three CBCL scales for the unadjusted models). Panel B: model A is the primary specification as reported; model B is the same specification restricted to the 3,095 participants with baseline waist-to-height ratio (6 fewer), the reference for model C; model C adds baseline waist-to-height ratio (*β* = 0.584 [0.547, 0.621]). The adversity coefficient fell by 47.8% from model B to model C and remained significant. *R^2^* = total model *R^2^*. *n* families: 2,783 (A) and 2,777 (B, C). CBCL = Child Behavior Checklist; FDR = false discovery rate; OLS = ordinary least squares.

***Table S15.*** *Baseline-Only, One-Per-Family, and Held-Out-Site Sensitivity Analyses*

| Specification | *N* train / held-out | Adversity features | Total SA held-out \|*r*\| | Retention, % [95% CI] | rsFC held-out \|*r*\| | Retention, % [95% CI] | Fluid β (*n*) | Crystallized β (*n*) |
| --- | --- | --- | --- | --- | --- | --- | --- | --- |
| Primary (reference) | 4,186 / 1,790 | 104 | 0.225 | 100 | 0.207 | 100 | -0.132 (1,148) | -0.109 (1,188) |
| Baseline-only battery | 4,186 / 1,790 | 71 | 0.221 | 98.5 [95.9, 101.1] | 0.201 | 96.9 [93.1, 100.6] | -0.132 (1,148) | -0.112 (1,188) |
| One participant per family | 3,712 / 1,591 | 104 | 0.229 | 101.9 [96.2, 108.0] | 0.206 | 99.4 [92.1, 107.3] | -0.134 (1,010) | -0.111 (1,048) |
| Held-out sites (15 train / 6 test) | 4,373 / 1,603 | 104 | 0.297 | 132.0 [107.2, 167.0] | 0.209 | 100.5 [76.9, 131.6] | -0.209 (1,082) | -0.093 (1,093) |

**Note.** Each specification re-estimated the training-only residualization on its own training rows. Structural and functional associations are held-out canonical |*r*| from two-view sparse models (L1 = 0.4): adversity view vs total cortical SA (single feature) and vs the 78-feature rsFC view. Retention = held-out |*r*| as a percentage of the primary held-out |*r*|, with 95% percentile CIs from 5,000 family bootstraps (training weights fixed); for held-out sites, site-resampled intervals were [99.5, 176.5] (SA) and [65.1, 148.8] (rsFC), and each held-out feature was centered within its site after the training age, sex, and motion coefficients were applied. The total-SA retention above 100% for held-out sites reflects a held-out |*r*| exceeding training |*r*|, not a stronger association. Baseline-only removes the 25 Life Events (year 1) and 8 neglect (year 3) items. Fluid and crystallized *β* = standardized Year-6 NIH Toolbox (uncorrected) coefficients for each specification’s own refitted adversity variate in its held-out participants, with the primary covariates; all *p* < .001. These analyses did not refit the five-component joint model. rsFC = resting-state functional connectivity; SA = surface area.

***Table S16.*** *Primary Model Versus Site-Unresidualized Adversity View and L1 = 0.7 Penalty*

| Quantity | Primary (L1 = 0.4) | Site not residualized from adversity view | L1 = 0.7 |
| --- | --- | --- | --- |
| *Dimensionality* | | | |
| Adversity-brain statistic, training | 0.0128 | 0.0106 | 0.0139 |
| Permutation *p* | <.001 | <.001 | <.001 |
| Adversity-brain statistic, held-out | 0.0125 | 0.0096 | 0.0126 |
| Permutation *p* | <.001 | <.001 | <.001 |
| Components replicating (of 5) | 1 | 1 | 1 |
| *Held-out \|r\| with the adversity variate (permutation p)* | | | |
| Surface area | 0.209 (<.001) | 0.178 (<.001) | 0.224 (<.001) |
| Resting-state connectivity | 0.190 (<.001) | 0.177 (<.001) | 0.173 (<.001) |
| Cortical thickness | 0.000 (.993) | 0.009 (.663) | 0.018 (.398) |
| Fractional anisotropy | 0.030 (.293) | 0.025 (.349) | 0.050 (.102) |
| MID task | 0.034 (.094) | 0.021 (.221) | 0.028 (.198) |
| Stop-signal task | 0.052 (.049) | 0.040 (.111) | 0.053 (.049) |
| Emotional n-back | 0.055 (.079) | 0.040 (.245) | 0.038 (.161) |
| Retention, SA, % [95% CI] | 100 | 85.0 [75.1, 95.0] | 107.2 [99.3, 116.3] |
| Retention, rsFC, % [95% CI] | 100 | 93.2 [82.9, 104.4] | 90.7 [71.8, 112.5] |
| *Composition relative to the primary solution* | | | |
| Non-zero weights, adversity (of 104) | 39 | 39 | 103 |
| Non-zero weights, SA (of 68) | 22 | 23 | 47 |
| Non-zero weights, rsFC (of 78) | 21 | 20 | 54 |
| *r* (weights), adversity | 1.000 | 0.992 | 0.903 |
| *r* (weights), SA | 1.000 | 1.000 | 0.841 |
| *r* (weights), rsFC | 1.000 | 0.998 | 0.397 |
| *r* (structural loadings), adversity | 1.000 | 0.986 | 0.979 |
| *r* (structural loadings), SA | 1.000 | 1.000 | 0.978 |
| *r* (structural loadings), rsFC | 1.000 | 0.999 | 0.845 |
| *r* (variate scores), adversity | 1.000 | 0.905 | 0.958 |
| *r* (variate scores), SA | 1.000 | 1.000 | 0.978 |
| *r* (variate scores), rsFC | 1.000 | 0.999 | 0.678 |
| Stable features (≥ 0.90), adversity | 19 | 18 | Not run |
| Stable features (≥ 0.90), SA | 12 | 11 | Not run |
| Stable features (≥ 0.90), rsFC | 10 | 6 | Not run |
| Ten largest adversity loadings | 5 neighborhood, 5 household SES | 5 neighborhood, 5 household SES | 5 neighborhood, 5 household SES |
| Site *R*² of training adversity variate (%) | 0.0 | 17.3 | 0.0 |

**Note.** Joint eight-view SGCCA, component 1. The site analysis residualized the adversity view on age and sex only (imaging views unchanged); the penalty analysis used the primary blocks at L1 = 0.7, the cross-validation optimum. Adversity-brain statistic = mean squared correlation of the adversity variate with the seven brain variates; permutation *p* from 10,000 draws for component 1 (minimum attainable *p* = .0001); components 2-5 were tested with 10,000 draws on deflated views and none replicated. Retention = sensitivity held-out |*r*| as a percentage of the primary held-out |*r*|, 95% CI from 5,000 held-out-family bootstraps. Weight, loading, and variate correlations compare each solution with the primary one across features (weights, training structural loadings) or participants (variate scores, all participants). Stability selection (10,000 subsamples) was run for the site analysis only. SES = socioeconomic status; SGCCA = sparse generalized canonical correlation analysis; rsFC = resting-state functional connectivity; SA = surface area; MID = monetary incentive delay.

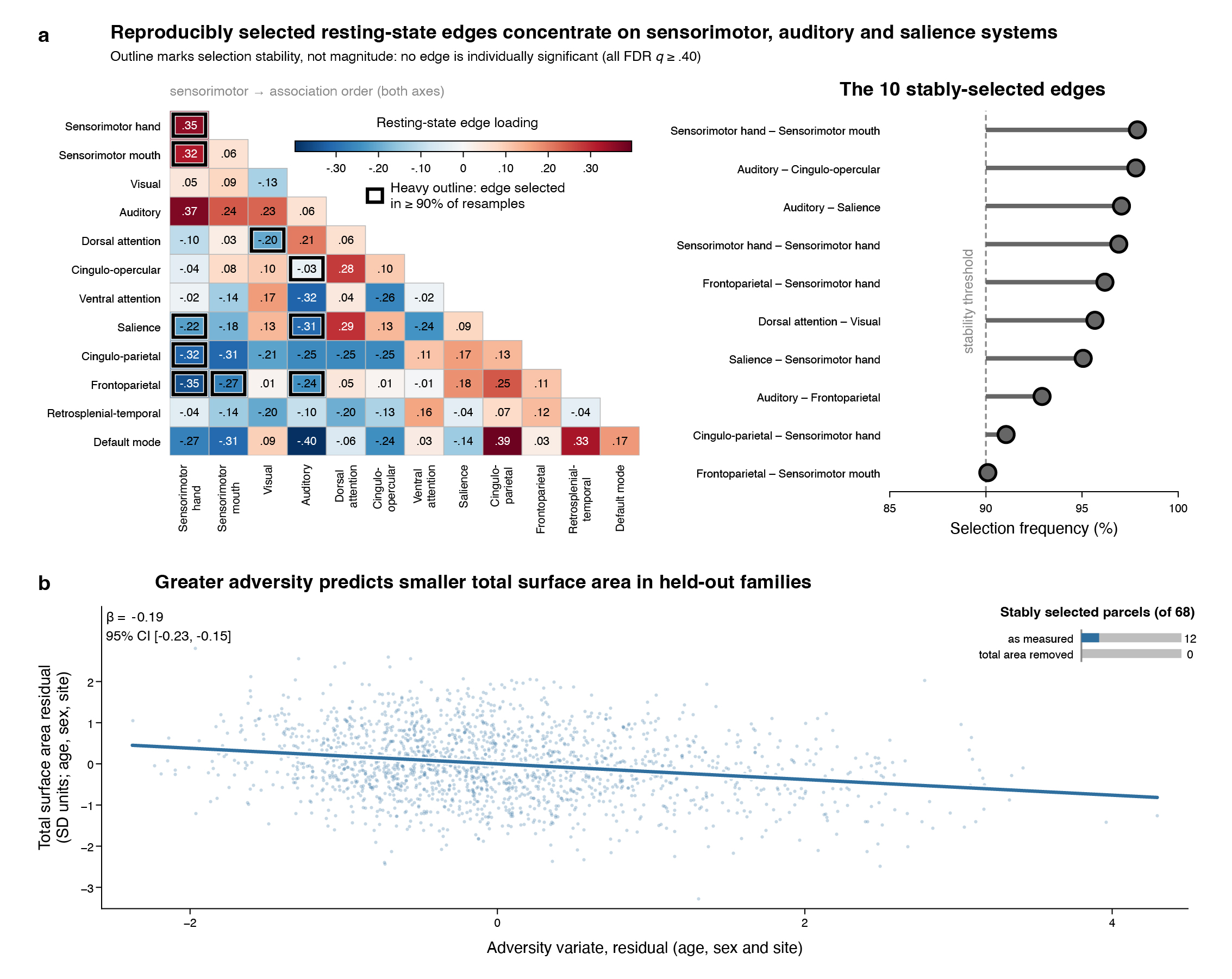

### Supplementary Figures

***Figure S1.*** *Resting-State Network Pair Loadings and the Global Surface Area Association*

**Note***.* (a) Left, structural loadings of all 78 rsFC network-pair features on the rsFC variate, shown as the lower triangle and diagonal of the 12 x 12 Gordon community matrix (each pair is one feature). Communities are ordered *a priori* from sensorimotor to association systems. The scale is diverging and centered at zero, and every cell prints its value. Heavy outlines mark the ten features selected in at least 90% of 10,000 subsamples. Right, the same ten features ordered by selection frequency; the dashed line marks the 90% threshold. Outlines denote selection reliability, not significance: no individual loading survived FDR correction (all *q* ≥ .405). Full values are in Table S9. (b) Added variable plot of total cortical SA against the adversity variate in the 1,790 held-out participants, with both residualized on age, sex, and site, so the fitted slope equals the standardized coefficient (*β* = -0.19, family-clustered 95% CI [-0.23, -0.15]). Inset, the number of the 68 SA parcels stably selected in the two-view model as measured (12) and after total SA was residualized from each parcel (0), over the same 2,000 subsamples. The association is cross-sectional and between families. SA = surface area; rsFC = resting-state functional connectivity; FDR = false discovery rate.
